# Natural variation in insect egg-induced cell death uncovers a role for L-type LECTIN RECEPTOR KINASE-I.1 in Arabidopsis

**DOI:** 10.1101/2020.05.29.122846

**Authors:** Raphaël Groux, Caroline Gouhier-Darimont, Envel Kerdaffrec, Philippe Reymond

## Abstract

In *Arabidopsis thaliana*, a hypersensitive-like response (HR-like) is triggered underneath the eggs of the large white butterfly *Pieris brassicae*, and this response is dependent on salicylic acid (SA) accumulation and signaling. Previous reports indicate that the clade I L-type lectin receptor kinase LecRK-I.8 is involved in early steps of egg recognition. A genome-wide association study (GWAS) was used to better characterize the genetic structure of HR-like and discover loci that contribute to this response. We report here the identification of LecRK-I.1, a close homolog of LecRK-I.8, and show that two main haplotypes that explain part of the variation in HR-like segregate amongst natural Arabidopsis accessions. In addition, signatures of balancing selection at this locus suggest that it may be ecologically important. Disruption of LecRK-I.1 resulted in decreased HR-like and SA signaling, indicating that this protein is important for the observed responses. Furthermore, we provide evidence that LecRK-I.1 functions in the same signaling pathway as LecRK-I.8. Altogether, our results show that the response to eggs of *P. brassicae* is controlled by LecRKs that operate at various steps of the signaling pathway.

## Introduction

Although eggs of herbivorous insects deposited on plant leaves are immobile and inert structures, they represent a future threat when hatching larvae start to feed. Plants are able to detect the presence of eggs and respond by triggering direct and indirect defenses (Reymond, 2013; Hilker & Fatouros, 2015). Direct defenses such as tissue growth and localized cell death lead to reduced egg hatching, egg crushing by surrounding tissues, or egg desiccation and drop off (Shapiro & DeVay, 1987; Doss *et al.,* 2000; Garza *et al.,* 2001; Desurmont & Weston, 2011; Petzold-Maxwell *et al.,* 2011; Fatouros *et al.,* 2014). The production of ovicidal benzyl cyanide at the oviposition site was reported in *Oryza sativa* (Suzuki *et al.,* 1996; Yamasaki *et al.,* 2003). For indirect defenses, a release of volatiles or changes in leaf surface chemistry attract natural egg predators (Hilker & Meiners, 2006; Fatouros *et al.,* 2012). While these defense mechanisms impact egg mortality individually, some studies found that the co-induction of both direct and indirect defense strategies synergistically impact egg survival (Fatouros *et al.,* 2014). Even though such defenses can efficiently reduce insect pressure before damage occurs, one study indicates that these traits may get lost during domestication (Tamiru *et al.,* 2015), similar to what is observed for other defense-related traits (Chen *et al.,* 2015). Introgression of egg-killing traits in cultivated varieties has been reported in *O. sativa* (Suzuki *et al.,* 1996; Yamasaki *et al.,* 2003; Yang *et al.,* 2014), however it is still a mostly unexploited strategy due to a lack of mechanistic understanding of these responses at molecular and cellular levels (Reymond, 2013; Fatouros *et al.,* 2016).

Plants from the Brassicales and Solanales orders respond to oviposition through the induction of cell death (Petzold-Maxwell *et al.,* 2011; Fatouros *et al.,* 2014, 2016; Kalske *et al.,* 2014), a process that resembles the hypersensitive response (HR) triggered by certain adapted pathogens. Based on this intriguing similarity, this type of insect egg-triggered response is referred to as hypersensitive-like response (HR-like) (Reymond, 2013; Fatouros *et al.,* 2014). Recent progress in the study of HR-like in response to insect eggs revealed that reactive oxygen species (ROS) and salicylic acid (SA), two major signaling molecules regulating plant immunity, accumulate at oviposition sites (Little *et al.,* 2007; Bruessow *et al.,* 2010; Geuss *et al.,* 2017; Bittner *et al.,* 2017; Bonnet *et al.,* 2017). In *Arabidopsis thaliana*, cell death induction upon crude egg extract (EE) treatment was reduced in mutants impaired in SA biosynthesis and signaling (Gouhier-Darimont *et al.,* 2013). In *Brassica nigra*, only plants expressing HR-like symptoms displayed elevated *PR1* transcript levels, a widely used SA marker gene, again suggesting that SA accumulation is required for HR-like induction (Fatouros *et al.,* 2014). These results indicate that the response to insect eggs is similar to pathogen-triggered immunity (PTI), as demonstrated in Arabidopsis (Gouhier-Darimont *et al.,* 2013). However, knowledge of elicitor-receptor pairs involved in plant response to eggs is lacking. A recent report identified the Arabidopsis L-type LecRK-I.8 as an early signaling component of *Pieris brassicae* egg-induced responses, thereby constituting a potential candidate for the perception of egg-derived elicitors (Gouhier-Darimont *et al.,* 2019). Lectin-like receptor kinases (LecRKs) have been implicated in a myriad of immune-related processes such as PTI (Takahashi *et al.,* 2007), chitin perception (Miya *et al.,* 2007), e-(extracellular) ATP and eNAD^+^/eNADP^+^ perception (Choi *et al.,* 2014; Wang *et al.,* 2019; Pham *et al.,* 2020), bacterial short chain 3OH-FA perception (Kutschera *et al.,* 2019), and insect resistance (Gilardoni *et al.,* 2011; Liu *et al.,* 2015).

Although insect eggs trigger a PTI-like response in Arabidopsis, identification of early and late components is still needed. Transcriptome changes after *P. brassicae* oviposition and pathogen infection are similar but not identical, suggesting some level of specificity (Little *et al.,* 2007). In support of this hypothesis, ROS production following *P. brassicae* EE treatment was independent on the NADPH oxidases RBOHD/F (Gouhier-Darimont *et al.,* 2013), while this pathway is crucial for pathogen-triggered ROS accumulation (Torres *et al.,* 2002; Morales *et al.,* 2016). Moreover, a *lecrk-I.8* mutant plant did not show altered resistance to a bacterial pathogen (Gouhier-Darimont *et al.,* 2019), suggesting that LecRK-I.8 may be specifically involved in egg signaling responses. To get more molecular insight in insect-egg induced response, we used the existing natural variation in EE-triggered HR-like in Arabidopsis and performed a genome-wide association study (GWAS) using a set of 295 natural accessions. We found a peak of association on chromosome 3 encompassing the clade I *L-type LecRK-I.*1, a close homolog of *LecRK-I.8*. Subsequent experimental validation showed that LecRK-I.1 is specifically involved in the regulation of cell death following insect egg perception, hence providing a novel component of this immune response.

## Materials and Methods

### Plant materials and growth conditions

All experiments described here were carried out in *Arabidopsis thaliana*. Plants were grown in growth chambers in short day conditions (10 h light, 22°C, 65% relative humidity, 100 μmol m^−2^ s^−1^) and were 4 to 5 weeks old at the time of treatment. Larvae, eggs and butterflies of the Large White butterfly *Pieris brassicae* came from a population maintained on *Brassica oleracea* in a greenhouse (Reymond *et al.,* 2000).

Accessions used for GWAS mapping were obtained from the NASC stock center and are listed in Supporting Information Table S1. T-DNA insertion lines for *lecrk-I.1* (SALK_052123), *lecrk-I.2* (SAIL_847_F07), *lecrk-I.3* (SALK_087804C), *lecrk-I.4* (SALK_091901), *lecrk-I.5* (GABI_777H06), *lecrk-I.6* (GABI_353G10) and *lecrk-I.8* (SALK_066416) were described previously (Gouhier-Darimont *et al.,* 2019). Primers for genotyping insertion lines were designed with the SIGNAL T-DNA verification tool for all lines used in this study.

A dual sgRNA CRISPR Cas9 approach (Pauwels *et al.,* 2018) was used to create a 33 kb deletion on chromosome 3 between *LecRK-I.1* (At3g45330) and *LecRK-I.6* (At3g45440), generating the sextuple mutant *ccI.1-I.6*. sgRNAs specific for *LecRK-I.1* and *LecRK-I.6* were selected using CRISPR-P (http://crispr.hzau.edu.cn/CRISPR2/). Complementary oligos with 4 bp overhangs were annealed and inserted in the Gateway ENTRY sgRNA shuttle vectors (Pauwels *et al.,* 2018) pMR217 (L1-R5) for sgRNA-LecRK-I.1 and pMR218 (L5-L2) for sgRNA-LecRK-I.6 via a cut-ligation reaction with BbsI (New England Biolabs) and T4 DNA ligase (New England Biolabs), generating two sgRNA modules pMR217-sgRNA-LecRK-I.1 and pMR218-sgRNA-LecRK-I.6. Using a Gateway Multisite LR reaction (Thermofisher), the two sgRNA modules were combined with pDE-Cas9Km (Pauwels *et al.,* 2018) to get the final expression clone that was inserted in *Agrobacterium tumefaciens* strain GV3101 for transformation of Arabidopsis Col-0 by floral dipping. Seeds were selected on ½ MS with 50 μg/ml kanamycin under continuous day conditions for 12 days. Presence of a deletion was tested by PCR, using specific primers CC-LecRK-I.1-Rv and CC-LecRK-I.6-Fw flanking the deletion and generating a 465 bp fragment (Figure S1).

### Genome-wide association study and haplotype analysis

For GWAS analysis, a set of 295 accessions from the RegMap panel (Horton *et al.,* 2012) were used (Table S1). Pools of 30 accessions were phenotyped every week over 2 days. For each accession, 3 leaves from 3 to 6 plants were treated with egg-extract diluted 1:1 with deionized water leading to a total of 9 to 18 treated leaves. Treated plants were left in the growth chamber for an additional 5 days until phenotyping. After 5 days, treated leaves were removed with forceps, symptoms were scored and pictures were taken as described below.

Mapping was performed locally using custom-made R scripts (R-Core Development Team, 2005; Kerdaffrec *et al.*, 2016) with a linear mixed model that accounts for population structure by computing a population-wide kinship matrix. Full imputed genotypes for 2029 lines (Togninalli *et al.*, 2018) were used and only SNPs with a minor-allele frequency (MAF) > 0.05 were considered for analysis. After filtering, a total of 1’769’548 SNP were used for mapping. To correct for genome-wide multiple testing, a Bonferroni corrected threshold of significance was computed by dividing α = 0.05 by the number of SNPs used in the analysis. The percentage of variance explained by the top SNP (Chr. 3, 16633422) of *LecRK1-1* was calculated using the residual sum of squares (RSS) extracted from the GWA model.

Haplotype analysis was based on the most significantly associated SNPs (−log_10_ *P* > 4) using data extracted from the genotype matrix. Presence/absence of premature stop codons in sequenced accessions (Table S1) was recovered from the POLYMORPH1001 Variant browser (https://tools.1001genomes.org/polymorph/). Both sets of data were used to explore phenotypic data.

The population-wide cladogram was built by calculating the Euclidian distance between accessions from the kinship matrix, and clustering was subsequently performed using the “ward.d2” algorithm in R. Gene sequences in Fasta format from 125 accessions were obtained from SALK 1001 genome browser (http://signal.salk.edu/atg1001/3.0/gebrowser.php) and sequences were aligned using MAFFT for further analysis.

Sliding windows analyses of Tajima’s D, Fu and Li’s D / F on the LecRK-I.1 locus were performed using DnaSP (version 6.12.03) with sliding windows of 200 variant sites and a step size of 25 sites, and significance threshold was set according to a 95% confidence limit published before (Tajima, 1989; Fu & Li, 1993). Homologous gene sequences from *AtLecRK-I.1* in *Arabidopsis lyrata* were identified by BLAST, and the best hit was used as outgroup sequence for Fu and Li’s statistics. 50 kb sliding-window analysis of Tajima’s D was computed using VCF genotype file obtained from the 1001 Genome API (https://tools.1001genomes.org/api/index.html). Tajima’s D was computed on sliding windows of 200 bp with a step size of 25 bp using custom-made R scripts.

### Broad sense heritability and variance analysis

For each phenotype, broad sense heritability (*H^2^*) was calculated using the following equations: *H^2^* = V_g_/V_p_ and V_p_ = V_g_ + V_e_, where V_g_, V_p_ and V_e_ stand for genetic, phenotypic and environmental variance respectively. Since accessions are homozygous, V_e_ was estimated as the average intra-accession variance and V_p_ was the variance of the phenotype in the population.

### Oviposition and treatment with EE

For experiments with natural oviposition, plants were placed in a tent containing approximately 20 *P. brassicae* butterflies for a maximum of 8 h. Afterwards, plants were kept in a growth chamber in plastic boxes until hatching of the eggs. Control plants were kept in the same conditions without butterflies.

*P. brassicae* eggs were collected and crushed with a pestle in Eppendorf tubes. After centrifugation (15’000 g, 3 min), the supernatant (‘egg extract’, EE) was collected and stored at −20°C. Plants were 4-5 weeks old at the time of treatment. For each plant, each of two leaves were treated with 2 μl of EE. This amount corresponds to one egg batch of ca. 20 eggs. A total of four plants were used for each experiment. After the appropriate time, EE was gently removed with a scalpel blade and treated leaves were stored in liquid nitrogen. Untreated plants were used as controls.

For GWAS, a large amount of EE was prepared as described and aliquots were stored under N_2_ at −80°C in order to ensure homogenous treatments during the entire experiment.

### Symptom scoring

Symptoms were scored visually from the adaxial side of the leaves and were classified into the following categories: no symptom (leaf treated area is still lush green, score = 0), small chlorosis (<50% of the treated area, score = 1), large chlorosis (> 50%, score = 2), small necrosis (brown spots or transparent membrane on < 50% of the treated area, score = 3), large necrosis (> 50 %, score = 4) and spreading necrosis (necrosis not confined to the treated area, score = 5).

### Histochemical staining

For visualization of cell death, EE was gently removed and leaves were submerged in lactophenol trypan blue solution (5 ml of lactic acid, 10 ml of 50% glycerol, 1 mg of trypan blue (Sigma), and 5 ml of phenol) at 28°C for 2–3 h. After staining, leaves were destained for 10 min in boiling 95% ethanol. Microscope images were saved as TIFF files and processed for densitometric quantification with ImageJ (version 1.48).

### Salicylic acid quantifications

SA quantifications were performed using the bacterial biosensor *Acinetobacter sp*. ADPWH (Huang *et al.,* 2005; Huang *et al.,* 2006) according to (DeFraia *et al.,* 2008; Zvereva *et al.,* 2016). Briefly, 6 leaf discs (0.7 cm, ~20 mg) were ground in liquid nitrogen and extracted in 0.1M sodium acetate buffer (pH 5.6). Extracts were then centrifuged at 4°C for 15 min at 16’000 g. 50 μL of extract were incubated with 5 μL of β-Glucosidase from almonds (0.5 U/μl in acetate buffer, Sigma-Aldrich) during 90 min at 37°C to release SA from SA-glucoside (SAG). 20 μL of extract was then mixed with 60 μL of LB and 50 μL of a culture of *Acinetobacter* sp. ADPWH_lux (OD_600_ = 0.4), and incubated for 1 h at 37°C. Finally, luminescence was integrated using a 485 ± 10 nm filter for 1 s. A SA standard curve diluted in untreated *sid2-1* extract amounts ranging from 0 to 60 ng was read in parallel to allow quantification. SA amounts in samples were estimated by fitting a 3^rd^ order polynomial regression on the standards.

### Gene expression analysis

Analysis of gene expression by quantitative real-time PCR was described previously (Bruessow *et al.*, 2010; Gouhier-Darimont *et al.*, 2013). Briefly, tissue samples were ground in liquid nitrogen, and total RNA was extracted using ReliaPrep™ RNA Tissue Miniprep (Promega) according to the manufacturer’s instructions. Afterwards, cDNA was synthesized from 500 ng of total RNA using M-MLV reverse transcriptase (ThermoFisher) and subsequently diluted eightfold with water. Reactions were performed using Brilliant III Fast SYBR-Green QPCR Master Mix (Agilent) on a QuantStudio 3 real-time PCR instrument (ThermoFisher). Values were normalized to the housekeeping gene *SAND* (At2g28390). Primer efficiencies were evaluated by five-step dilution regression. For each experiment, three biological replicates were analyzed. A list of all primers used in experiments can be found in Table S2.

Publically available RNA-seq data (GSE80744) from 728 accessions were recovered from the Gene Expression Omnibus repository (www.ncbi.nlm.nih.gov/geo/GEO). Expression data were then filtered to extract only values for accessions used in the study.

### Statistical analyses

GWAS and subsequent analysis of the data obtained was performed with R software version 3.6. For boxplots, the thick line indicates the median, box edges represent 1st and 3rd quartile respectively, whiskers cover 1.5 times the interquartile space and dots represent extreme values. Boxplot width is proportional to the number of sample. When displayed, notches indicate an approximate confidence interval for median values. All other analyses using mutant lines were analyzed using GraphPad Prism 8.0.1.

## Results

### Arabidopsis accessions display natural variation in response to P. brassicae eggs

We observed that Arabidopsis accessions display varying degrees of HR-like symptoms after 5 days of treatment with *P. brassicae* egg extract (EE) (Fig. 1a), corresponding to the hatching time of real eggs. The existence of natural variation for this defense-related trait suggests that it is under genetic control, consistent with previous reports (Gouhier-Darimont *et al.,* 2013, 2019). While most leaves exhibited some degree of chlorosis, symptoms usually ranged from no visible symptom to the formation of large patches of cell death (Fig. 1b). Moreover, symptoms were most of the time restricted to the area of egg application, but could exceptionally grow larger for both chlorosis and necrosis. However, because of the low frequency of such events, we did not quantify those separately. This scoring scheme allows an easy quantification of the HR-like (Fig. 1c). To verify that application of *P. brassicae* EE mimics HR-like responses induced by real eggs, we tested whether naturally oviposited eggs could also induce some degree of variation in HR-like responses. In several accessions, we observed the formation of chlorotic or necrotic tissue localized around and underneath the egg clutches after 5 days (Fig. 1d) and this correlated with the severity of symptoms observed after EE treatment. This response was however more variable, suggesting that EE treatment might be more consistent than real eggs where the protective shell may slow down the release of elicitors. We thus set conditions for a GWAS using EE treatment and performed initial tests on three selected accessions, using Col-0 as a control. Based on these experiments, we used short-day grown plants (10L:14D), treated three leaves per plant with EE diluted 1:1 with deionized water, and scored symptoms after five days (Fig. S2a-c). Because such experiment requires a large amount of EE, this allowed to maximize the overall number of plant treated per accession and per block, thereby improving power, even though the use of undiluted EE and/or long days conditions tend to increase symptom strength (Fig. S2a-c).

**Fig. 1.**
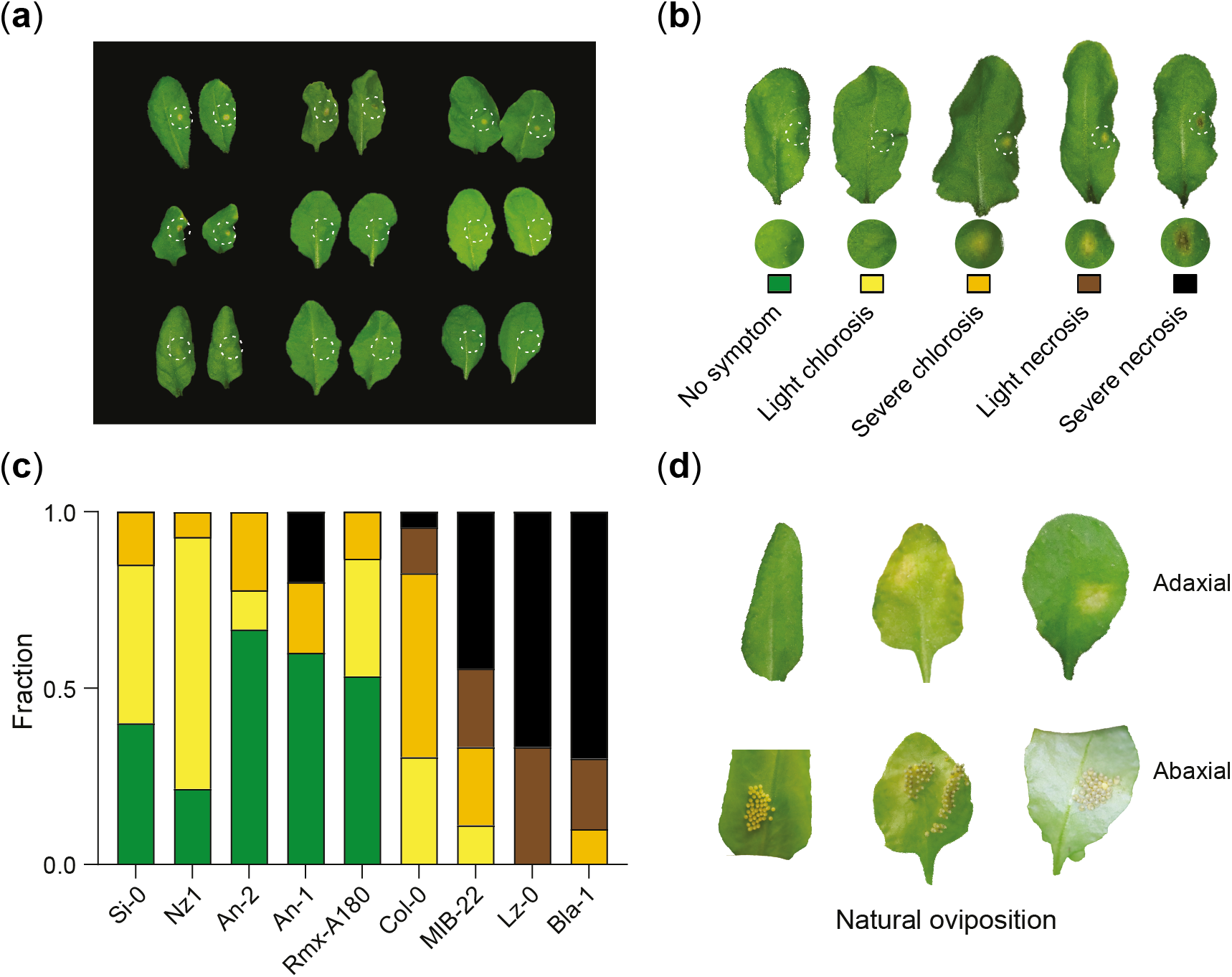
Insect egg-induced HR-like varies among natural Arabidopsis accessions. (a) Representative picture of accessions (two leaves per accession) displaying varying phenotypes after treatment with *P. brassicae* egg extract. (b) Representative pictures of symptoms used for scoring in experiments. (c) Proportion of each symptom class developed by some accessions shown in panel (a) as visualized from the adaxial side. (d) Variation in HR-like triggered after natural oviposition by *P. brassicae* butterflies. In all experiments shown here, duration of treatment was 5 days as this corresponds to the hatching time of naturally oviposited *P. brassicae* eggs. The dashed circles indicate the site of treatment on the abaxial side of the leaf.

### Identification of a genetic locus associated with P. brassicae egg-induced symptoms

Broad sense heritability (*H^2^*), which estimates the amount of phenotypic variance that is genetically encoded, was high for symptom score (*H^2^* = 0.56). This finding is consistent with previously reported *H^2^* values for defense phenotypes (Yang *et al.*, 2017). GWAS using 295 accessions and a fully imputed genotype matrix (Togninalli *et al.*, 2018) revealed the existence of two loci associated with the degree of cell death induced by EE (Fig. 2a). One peak of association was found on chromosome 2 (−log_10_ *P* = 15.96) and constitutes the basis for another study (Groux *et al.*, in preparation). Besides this peak, we observed the existence of another significantly associated marker (−log_10_ *P* = 7.59, position 16633422) on chromosome 3 (Fig. 2a), which explains a total of 9.56% of the phenotypic variance. By taking a closer look at this genomic region, we found that this marker is located in the coding sequence of the *L-TYPE LECTIN RECEPTOR KINASE-I.1* (At3g45330) (Fig. 2b). Interestingly, this genomic locus encompasses 5 other closely related L-type clade I *LecRK* genes, *LecRK-I.2* to *LecRK-I.6*. We found that high linkage disequilibrium (LD) with other surrounding markers was only observed for SNPs found in the gene sequence of *LecRK-I.1* (Fig. 2b), suggesting that this gene may be causal for the differences in HR-like symptoms observed. We then determined whether this locus was previously identified in other publicly available GWAS. According to the araGWAS database (Togninalli *et al.,* 2018), markers at the *LecRK-I.1* locus are not associated with any of the defense-related or developmental phenotypes available, suggesting that it may play a specific role during *P. brassicae* egg-induced response (Fig. S3).

**Fig. 2.**
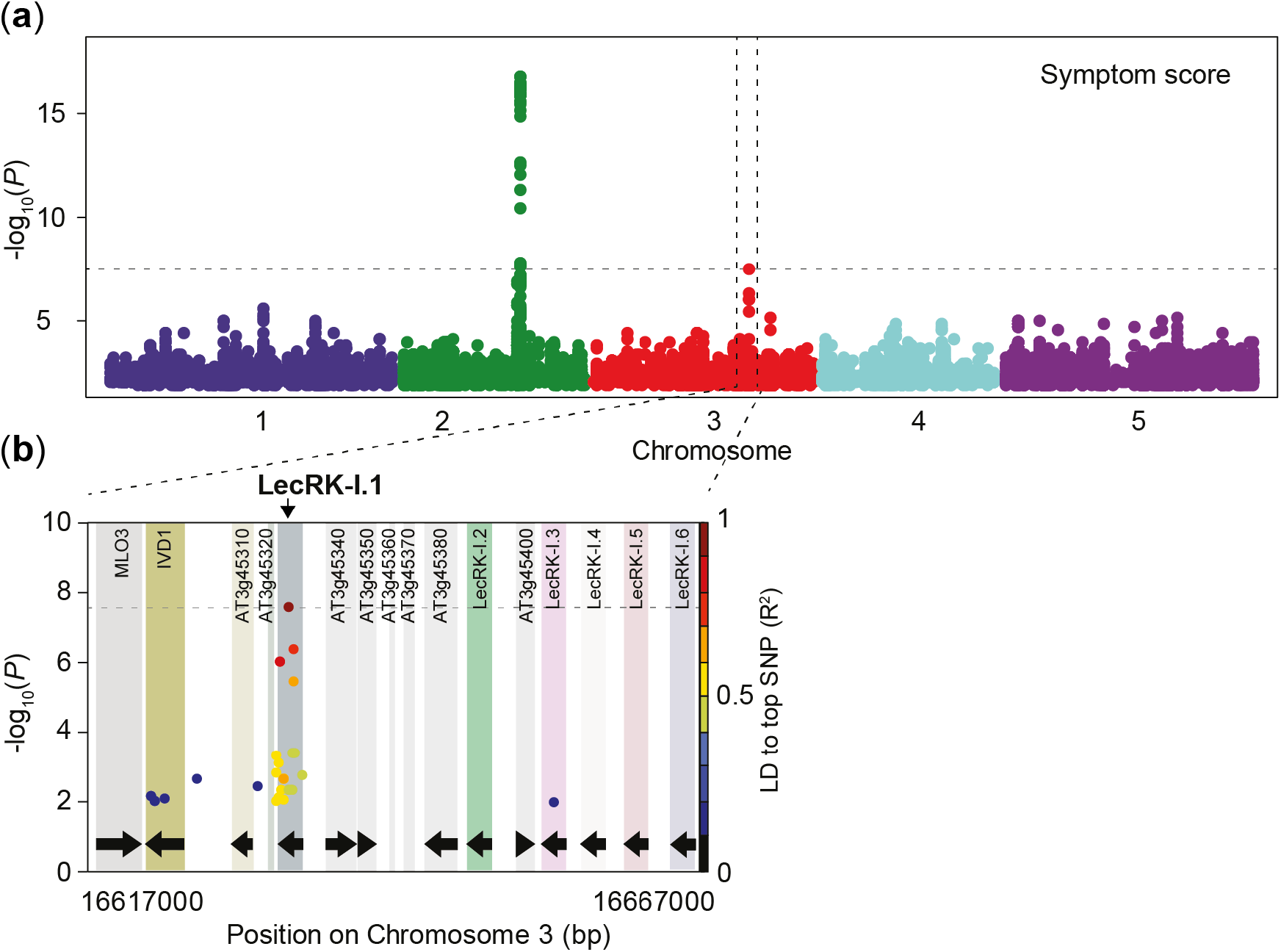
Genome-wide mapping of insect-egg induced HR-like symptoms. (a) Manhattan plot of the GWAS for symptom score after 5 days of *P. brassicae* EE treatment using a linear mixed-model. Full imputed genotypes (1’769’548 SNPs) for all 295 accessions were used for mapping. Chromosomes are displayed in different colors and the dashed line indicates a significance level of 0.05 after Bonferroni correction for multiple testing. (b) Local association plot of a 50 kb region surrounding the most significant marker. The x-axis represents the genomic position on chromosome 3 and color boxes indicate genes. LD to the most significant SNP (SNP3) is indicated by a color scale.

### LecRK-I.1 is involved in EE-induced cell death

In order to test whether clade I LecRKs found in this region could be involved in the response to *P. brassicae* EE, we tested whether the lack of a single LecRK using T-DNA insertion lines leads to alterations in different egg-induced responses. Additionally, we used CRISPR-Cas9 technology to delete the entire cluster of *LecRK-I.1* to *LecRK-I.6* (hereafter named *ccI.1-I.6*) in order to explore whether redundancy among these genes exists (Supporting Information Fig. S1). Symptom score was found to be significantly reduced in *lecrk-I.1* mutant but not in knock-out lines from surrounding homologs (Fig. 3a). Consistently, deletion of the gene cluster lead to symptom reduction similar to that observed in *lecrk-I*. We then quantified cell death at the site of EE application by trypan blue staining and observed a similar pattern (Fig. 3b). These results clearly show that *LecRK-I.1*, but not other homologs from the genomic cluster, is involved in the induction of HR-like in response to EE treatment.

**Fig. 3.**
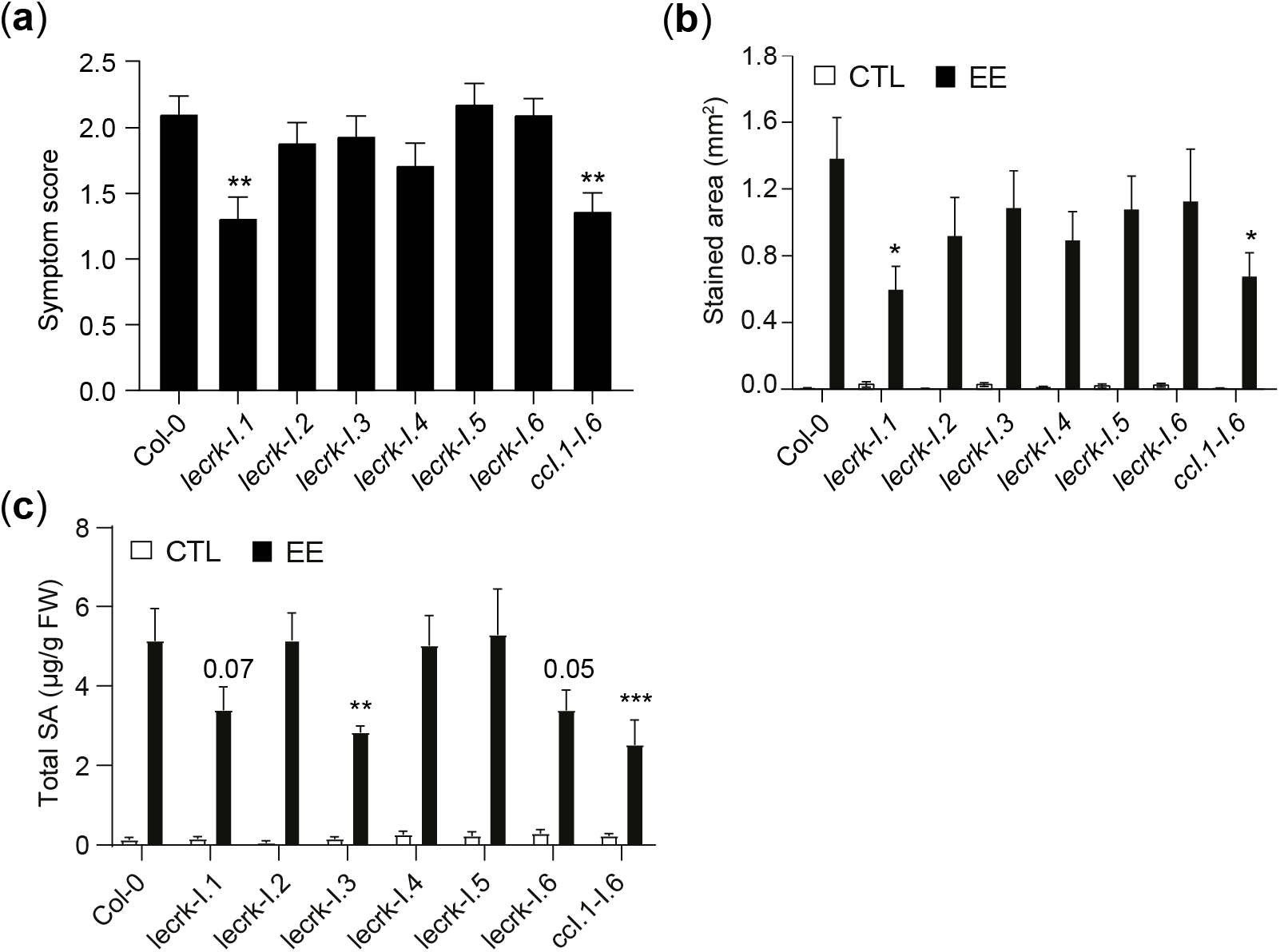
LecRK-I.1 plays a role in the induction of HR-like symptoms following insect egg perception. (a) Average symptom scores as visualized from the adaxial side after 5 days of treatment with EE. Means ± SE from three independent experiments are shown (n= 12-23 for each experiment). (b) Cell death as quantified by trypan blue staining after 3 days of EE treatment. Untreated leaves were used as controls. Means ± SE from 8-20 leaves are shown. This experiment was repeated once with similar results. (c) Total salicylic acid (SA + SAG) after 3 days of EE treatment. Measurements were done on 4 plants per treatment using a bacterial biosensor, untreated plants were used as control. Means ± SE of three independent experiments are shown (n= 12). Stars indicate significant differences with Col-0 (ANOVA followed by Dunnet’s test). *, *P*<0.05; **, *P*<0.01; ***, *P*<0.001.

Because altered SA signaling in the *lecrk-I.1* mutant could account for the reduction in cell death, we next quantified SA levels in plant treated with EE for 3 days. Total SA levels were reduced in *lecrk-I.1* after 3 days of treatment compared to wild-type plants, however this trend was not statistically significant (Fig. 3c). Interestingly, we observed a similar result in the *lecrk-I.6* mutant and a significant reduction in *lecrk-I.3*, indicating that multiple LecRKs may regulate SA accumulation. Removal of all six *LecRKs* in the *ccI.1-I.6* line resulted in significantly reduced SA levels after EE treatment (Fig. 3c), further supporting that one or more clade I LecRKs may participate in EE-induced SA accumulation. Altogether, these results demonstrate a role for LecRK-I.1 in the regulation of cell death after insect egg perception, while different clade I LecRKs regulate SA levels during this response.

### LecRK-I.1 haplotypes correlate with egg-induced symptoms

The finding that natural variation in *LecRK-I.1* is associated with the inducibility of HR-like responses among Arabidopsis accessions indicates that variants of this gene may alter protein function(s) or gene expression. As mentioned, we observed that a single marker reached genome-wide significance (SNP3; Chr. 3, 16633422) while four other SNPs had intermediate *P*-values (−log_10_ *P* > 4), all within the coding sequence of *LecRK-I.1* (Fig. 4a,b). Interestingly, three out of five SNPs result in amino acid changes: one (SNP5) in the putative carbohydrate-binding lectin-like domain (I228R), and two in the kinase domain (R356K, SNP3; I602V, SNP1), while the two other associated markers lead to silent mutations (Fig. 4b). Additionally, linkage analysis revealed that all markers are in moderate to strong LD with SNP3 (Fig. 4a). Since we do not know which marker might be causal, we considered haplotypes defined by SNP1-5. We next determined that at least 5 haplotypes are present in the population used for GWAS mapping (Fig. 4c), with two of them being present at rather high frequency compared to the other three (H2, 30.51%; H5, 58.64%; H1, H3 and H4 <7%). We observed that the distribution of symptom score was not different between haplotypes having variants in the lectin domain, suggesting that they might not impact this response. However, non Col-0 alleles for markers in the kinase and lectin domains together are associated with significantly lower symptoms (H2 vs H5, Fig. 4c). Together with the previous finding, these results potentially indicate that differential kinase function may underlie natural variation in EE-induced cell death. To explore this possibility, we examined whether these variants are found in functional subdomains of the kinase. Based on annotation from the UNIPROT database, we found that R356K lies in the predicted ATP binding site. We then performed homology-based modelling using SWISSMODEL (https://swissmodel.expasy.org) to predict the 3D topology of the LecRK-I.1 kinase domain in order to assess whether this amino acid change could potentially alter kinase function. We found that even though R356 is indeed proximal to the ATP binding site, the residue’s side chain was not within 4Å of the ATP molecule and is exposed to the solvent in the model obtained (P. Jimenez Sandoval, results not shown). It is thus unlikely to interact with ATP within the active site. Moreover, we found that this position does not seem to be conserved in the structure of immunity-related plant kinases such as BAK1, and in all cases side chains were exposed to the solvent. Finally, the predicted amino acid change (R to K) does not lead to significant steric or charge modification at this site. This was also true for the other kinase variant, and the lectin variants appeared to be outside of the canonical lectin-binding or multimerization interfaces and are also solvent exposed (P. Jimenez Sandoval, results not shown). These results suggest that the identified variant may not be involved in canonical kinase or receptor function based on knowledge obtained from homologous proteins. We also determined whether multiple independent frameshifts and/or premature stop codon may be present in natural variants at this locus, potentially leading to the production of a truncated protein. We found that 18% of the sequenced accessions (23/125) possessed such variants (Fig. S4a), yet this does not correlate with any difference in symptom distribution independently of the haplotype considered (Fig. S4b). Moreover, premature STOP codons occurred within the first 20 first amino acid of the sequence. This suggests that these frameshifts and premature stops may not necessarily lead to the production of non-functional proteins, possibly through the existence of alternative start sites. Collectively, our results point to a role for variation in the *LecRK-I.1* gene sequence in modulating HR-like symptoms triggered by *P. brassicae* EE. However, the link between variation in the coding of this gene and the resulting phenotype is still unclear.

**Fig. 4.**
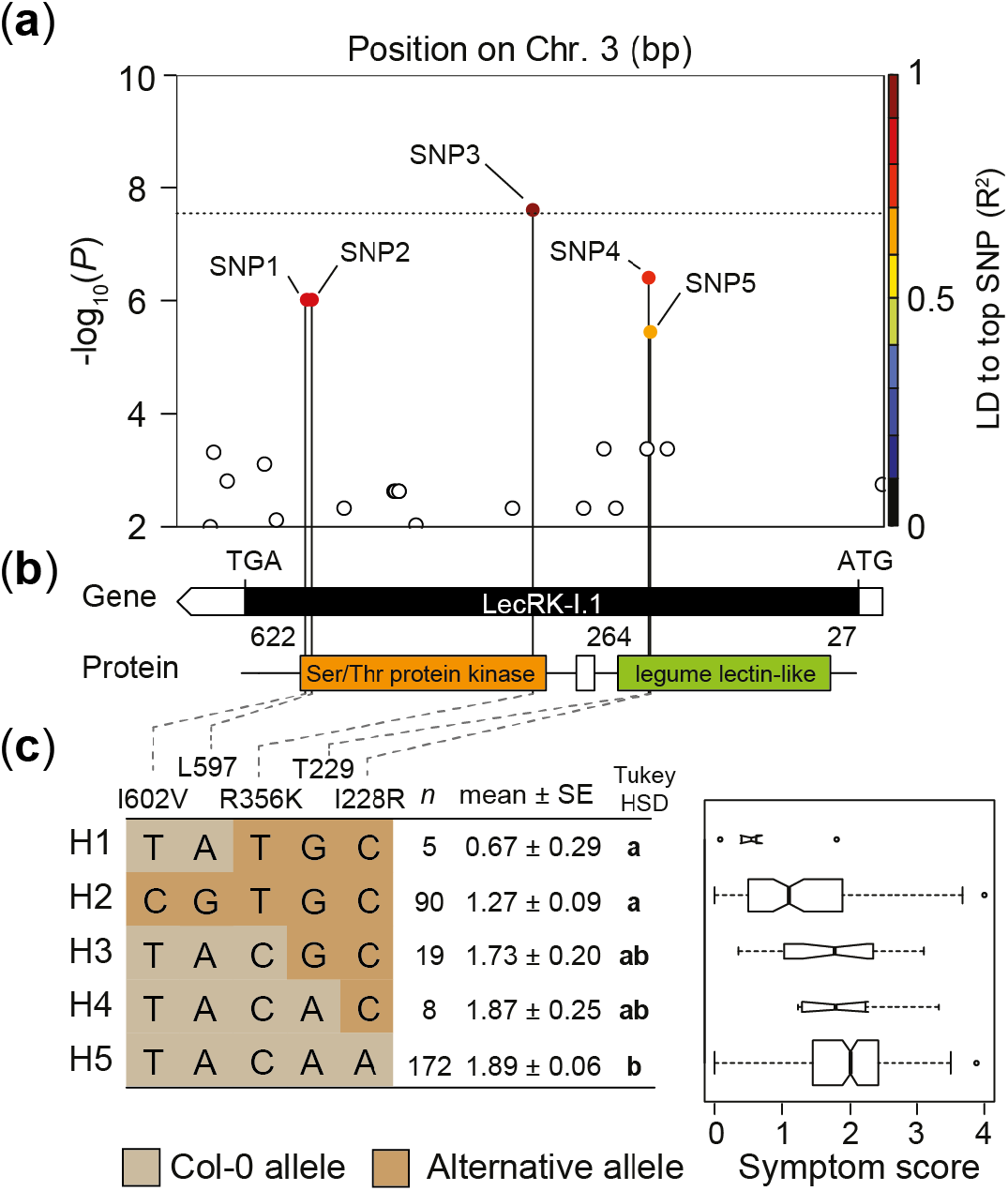
Local association and haplotype analysis of the *LecRK-I.1* locus. (a) Local association plot of the *LecRK-I.1* locus. The x-axis represents the genomic position on chromosome 3. LD of markers (−log_10_ *P*>4) to the most significant SNP (SNP3) is indicated by a color scale. The dashed line indicates the Bonferroni corrected significance threshold at α=0.05. (b) Gene and protein domain organization according to UNIPROT. (c) Haplotype analysis using the five most significant SNPs in the *LecRK-I.1* gene. Mean symptom score ± SE is shown and n indicates the number of accessions carrying each haplotype. Different letters indicate significant difference at *P*<0.05 (ANOVA, followed by Tukey’s HSD for multiple comparison). The coordinates on chromosome 3 of SNP1-5 displayed in panel (a) are the following: SNP1: 16632685; SNP2: 16632698; SNP3: 16633422; SNP4: 16633802; SNP5: 16633802.

### LecRK-I.1 expression is haplotype-specific but does not correlate with symptom severity

The absence of any obvious impact of the variation identified within the *LecRK-I.1* coding sequence may indicate that an alternative process is involved. We determined whether differences in gene expression may correlate with haplotype identity and HR-like symptoms by measuring *LecRK-I.1* transcript levels following EE treatment in 40 accessions with high or low symptoms. Gene expression was highly variable in the accessions surveyed, and expression was barely detectable in certain lines (Fig. S5). We found that the two most frequent haplotypes were not associated with differential *LecRK-I.1* expression after EE treatment, although one of the minor haplotype displayed comparatively lower transcript levels (Fig 4). Despite this haplotype-specific expression pattern, we found no correlation between symptom score and *LecRK-I.1* expression levels in the surveyed accessions (Fig. 5a,b). Analysis of publicly available RNA-seq data from different accessions (see methods) allowed for a similar analysis using SNP3 to split the data. Consistent with our experiments, none of the alleles at this position was associated with altered *LecRK-I.1* transcript levels (Fig. S6). Altogether these results suggest that variation in *LecRK-I.1* expression does not account for the variation in HR-like observed between accessions.

**Fig. 5.**
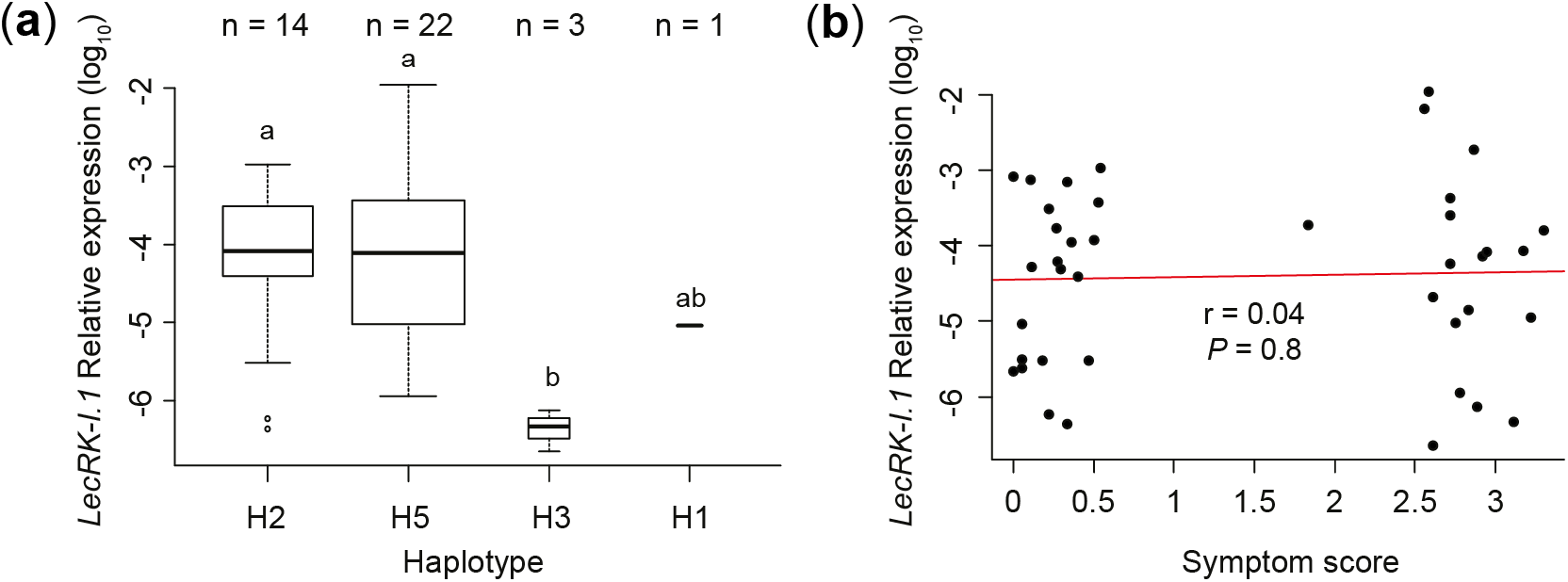
Natural *LecRK-I.1* haplotypes are not differentially expressed upon EE treatment and expression does not correlate with symptom score. (a) *LecRK-I.1* expression in 40 different accessions with low and high symptoms after 72 h of EE treatment. Transcript levels were plotted according to haplotypes defined in Fig. 4. Different letters indicate significant difference at *P*<0.05 (ANOVA, followed by Tukey’s HSD for multiple comparison) (b) *LecRK-I.1* expression in accessions with low or high symptoms does not correlate with symptom scores of the respective accessions. Gene expression was monitored by qPCR and target gene transcript level was normalized to the reference gene *SAND*. Means of three technical replicates are shown. Expression data were corrected by adding half the smallest non-zero value in order to avoid zero values, and log_10_-transformed prior to analysis. ns, not significant (ANOVA, *P*>0.05). Pearson correlation coefficient (r) and corresponding P value (*P*) between *LecRK-I.1* transcript level and symptom score are shown.

### Signatures of selection at the LecRK-I.1 locus

Based on the haplotype analysis, we found that only five *LecRK-I.1* haplotypes segregate in the mapping population used in this study. To get a closer look, we calculated haplotype frequencies using SNP1-5 described earlier. As mentioned before, two haplotypes appear to be present at high frequencies (H2, 30.51%; H5, 58.64%), while the three remaining haplotypes are much less frequent (< 7%, Fig. 6a). This could indicate that haplotypes may be distributed in two subpopulations due to adaptation to a type of environment for example, or that selection is acting at this loci and is maintaining variation. To disentangle these hypotheses, we constructed a cladogram representing genetic distance by using the kinship matrix used during the GWA mapping and superimposed haplotype identity for each accession (Fig. 6b). While it does appear that some low frequency haplotypes could be specific to certain geographical locations, the two major haplotypes appear to have a very broad distribution in the entire Arabidopsis population used for mapping (Fig. S7). The absence of any obvious link with phylogenetical or geographical history suggests that selection may be acting at this locus. To test the hypothesis that the *LecRK-I.1* gene sequence does not evolve neutrally, we measured Tajima’s D statistic along the coding sequence by using a sliding window (Fig. 6c). Tajima’s D compares the observed frequency of variants to those expected for a similar sequence evolving neutrally (Tajima, 1989), thereby indicating any departure from neutrality. We found that Tajima’s D was mainly positive along the gene sequence of *LecRK-I.1*, and two short stretches were found to be significantly positive (Fig. 6c). This result is indicative that *LecRK-I.1* might not evolve neutrally. In addition, we found that nearly all genes surrounding *LecRK-I.1* displayed negative Tajima’s D values when computed over a 50 kb region (Fig. S8). Because Tajima’s D is sensitive to demographic history, we also performed a sliding window analysis of Fu and Li’s D and F statistics, which considers the species history by including an outgroup (*Arabidopsis lyrata*). We observed that the pattern and sign of Fu and Li’s statistics were very similar to Tajima’s D (Fig. 6c), further supporting the hypothesis that this locus is not evolving neutrally.

**Fig. 6.**
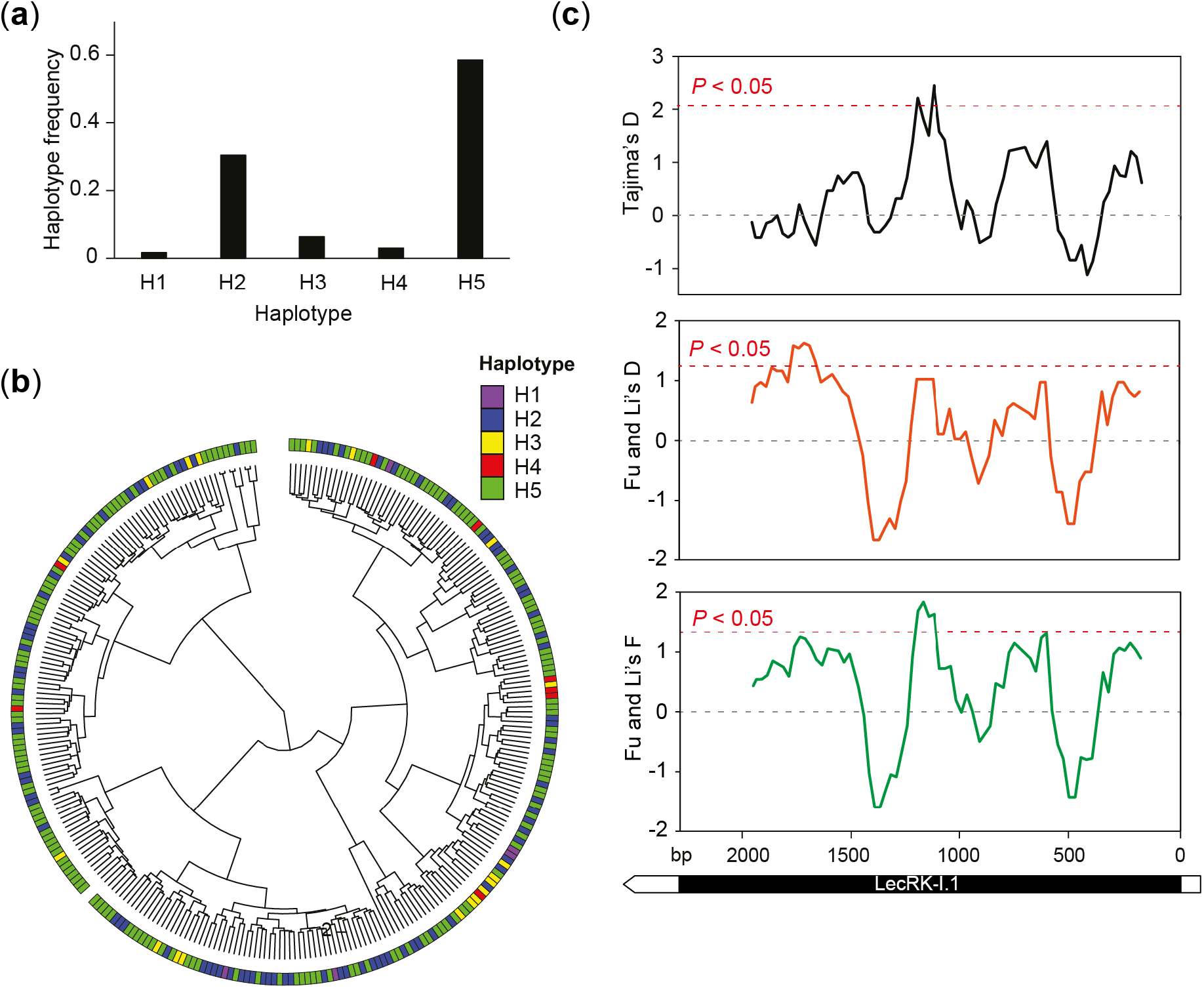
Signatures of selection at the *LecRK-I.1* locus. (a) Frequency of the different haplotypes in the accession panel used for GWAS mapping. (b) Cladogram constructed from a genome-wide kinship matrix of the 295 accessions used for GWAS. The outermost circle indicates the haplotype carried by a given accession. (c) Sliding window analysis of Tajima’s D, and Fu and Li’s D and F statistics along the *LecRK-I.1* coding region using a window size of 200 variant sites and a step size of 25 sites. A subset of 125 accessions with available full genome sequences was used for this analysis. The red dashed line indicates significance threshold at *P*<0.05. The gene structure of *LecRK-I.1* is shown below.

### LecRK-I.1 and LecRK-I.8 function in the same pathway

Based on the previously reported identification of LecRK-I.8, a close homolog of LecRK-I.1 (Bouwmeester and Govers, 2009; Hofberger *et al.,* 2015), as an early component of egg-triggered signaling (Gouhier-Darimont *et al.,* 2019), we wondered whether both genes may function in the same signaling cascade or if they are involved in parallel pathways. We crossed *lecrk-I.1* with *lecrk-I.8* and measured EE-induced responses in the single and double mutants. Both single mutants displayed a similar reduction of HR-like symptoms, *PR1* expression and SA accumulation after EE treatment. However, the double mutant did not show an enhanced reduction of these responses, strongly suggesting that LecRK-I.1 and LecRK-I.8 act in the same pathway (Fig. 7a-d).

**Fig. 7.**
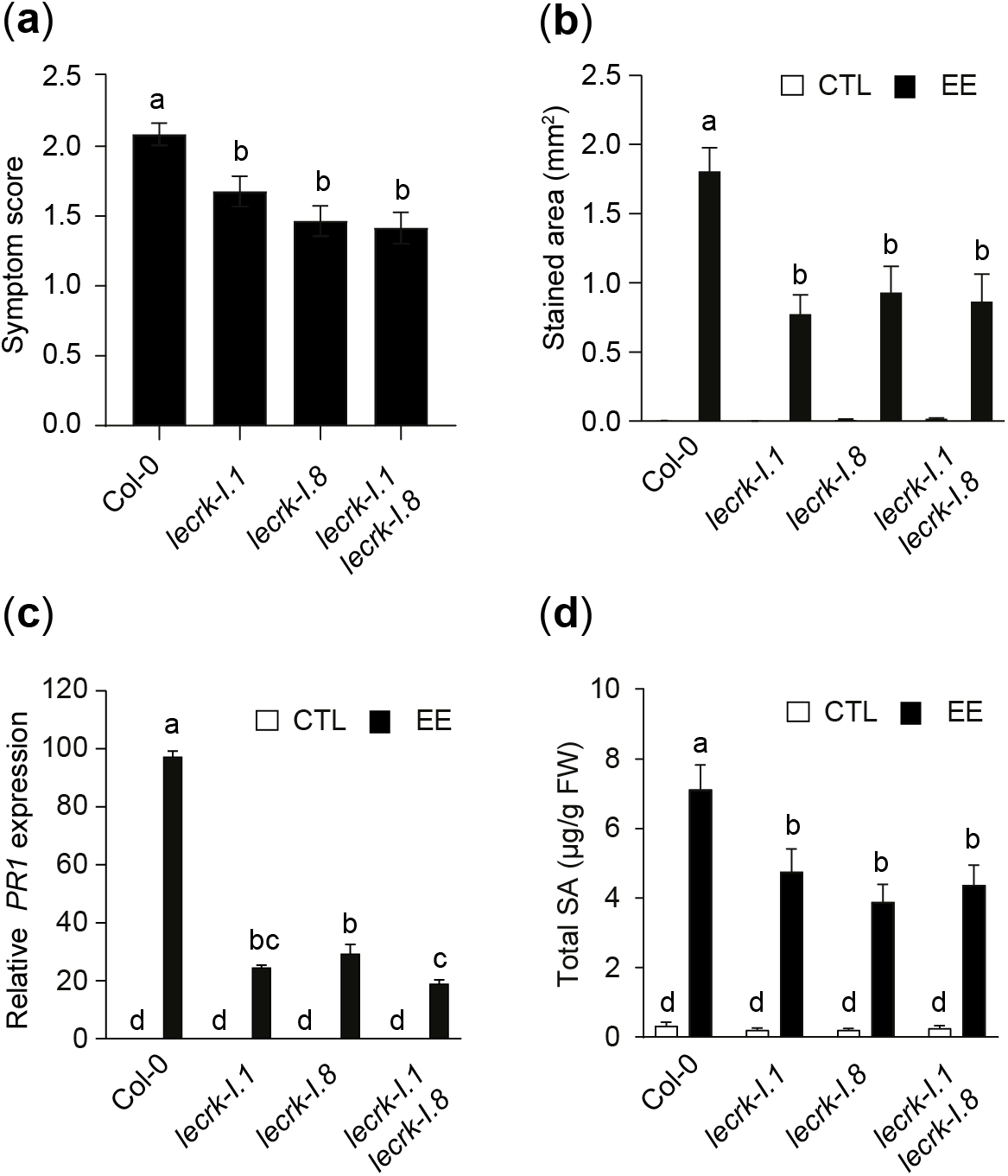
*LecRK-I.1* genetically interacts with *LecRK-I.8*. (a) Col-0, single and double *lecrk* mutants were treated with EE. Average symptom score was visualized from the adaxial side after 5 days of EE treatment. Mean ± SE from three independent experiment is shown (n=12-23 for each experiment). (b) Cell death as quantified by trypan blue staining after 3 days of EE treatment. Untreated leaves were used as controls. Means ± SE from 8-20 leaves are shown. This experiment was repeated once with similar results. (c) Expression of the marker gene *PR1* after 3 days of EE treatment. Transcript levels were monitored by qPCR and normalized to the reference gene *SAND*. Means ± SE of three technical replicates are shown. This experiment was repeated twice with similar results (d) Total salicylic acid (SA + SAG) after 3 days of EE treatment. Measurements were done on 4 plants per treatment using a bacterial biosensor, untreated plants were used as control. Means ± SE of three independent experiments are shown (n=12). Different letters indicate significant difference at *P*< 0.05 (ANOVA followed by Tukey’s HSD test for multiple comparison).

## Discussion

The recent description of LecRK-I.8 as an upstream regulator of insect egg-triggered signaling opened the exciting possibility that it could be involved in egg perception (Gouhier-Darimont *et al.,* 2019). However, several phenotypes including cell death were not completely abolished in the *lecrk-I.8* mutant, indicating a potential redundancy with other homologs or that LecRK-I.8 works as a modulator of the response. Indeed, expression all other clade I *L-type LecRKs* is also induced following EE treatment (Gouhier-Darimont *et al.,* 2019). We describe here the identification of LecRK-I.1 as a component of the insect egg-triggered signaling pathway in Arabidopsis that specifically affects the induction of HR-like during this interaction. We show that a knock-out mutant of *LecRK-I.1* leads to reduced HR-like symptoms and cell death following *P. brassicae* EE treatment, while SA accumulation is reduced. Moreover, double knock-out of both *LecRK-I.1* and *LecRK-I.8* did not result in a further reduction of HR-like symptoms, *PR1* expression and SA accumulation, suggesting that both genes function in the same transduction pathway or are part of the same complex. Other clade I *LecRK* mutants showed reduced SA levels and this was further confirmed by deleting all six genes clustered in the same locus. These results indicate that several clade I LecRKs are involved in the response to eggs and control different aspects of this pathway. It is noteworthy that only LecRK-I.1 and LecRK-I.8 affect the induction of cell death following egg perception, whereas only the *LecRK-I.1* locus displays natural variation.

Despite the absence of known ligands for LecRK-I.1 and most other clade I L-type LecRKs, it has been reported that LecRK-I.1 possesses a putative RGD-binding motif and that it may participate in plasma membrane-cell wall interactions (Gouget *et al.,* 2006). Whether insect eggs trigger changes in cell wall properties is not known, but epicuticular wax patterns were shown to be altered upon *P. brassicae* oviposition on Arabidopsis (Blenn *et al.,* 2012). In contrast to lectins, LecRKs show poorly conserved sugar-binding residues and it is therefore unlikely that they bind carbohydrates (Bouwmeester & Govers, 2009). However, the conserved hydrophobicity of the resulting cavity could recognize more hydrophobic ligands. Given the lipidic nature of egg-derived inducing compounds (Bruessow *et al.,* 2010; Gouhier-Darimont *et al.,* 2019), this would be consistent with a role for L-type LecRKs in egg perception. Recently, the G-type LecRK LORE was found to bind to medium chain 3-OH fatty acids that trigger LORE-dependent immunity (Kutschera *et al.,* 2019), further supporting the hypothesis that some LecRKs may recognize hydrophobic ligands from insect eggs. Alternatively, the identified LecRKs could also be involved in the perception of secondary signals such as DAMPs as suggested by the identification of LecRK-I.8 and LecRK-VI.2 as potential receptors for eNAD^+^ and eNADP^+^ (Wang *et al.,* 2017; Wang *et al.,* 2019), and LecRK-I.9 and LecRK-I.5 as eATP receptors (Choi *et al.*, 2014; Pham *et al.*, 2020). Here, we provide evidence that LecRK-I.1 and LecRK-I.8 function as components of the same pathway/complex. This suggests that LecRK-I.1 might serve as a secondary signaling hub that controls LecRK-I.8-dependent cell death. Whether LecRK-I.1 binds egg-derived elicitors alone or in association with LecRK-I.8 is an intriguing hypothesis that deserves further research. A more detailed evaluation of egg-triggered phenotypes in single and double mutants will help address this hypothesis.

We found that natural variation in the *LecRK-I.1* gene sequence was associated with HR-like symptoms, consistent with the phenotype observed in the *lecrk-I.1* mutant. Two main haplotypes of this gene segregate at the population level and we found that one variant, in particular (SNP3), located in the kinase domain was significantly associated with HR-like symptoms. Evaluation of *LecRK-I.1* expression in accessions indicates that transcript levels are not associated with symptom severity, therefore suggesting that functional variation in the LecRK-I.1 protein sequence modulates HR-like. Unfortunately, our analysis suggests that none of the identified variants would significantly affect the lectin binding pocket or the kinase domain. *In vitro* kinase assay using different variants of the kinase domain may help address this question directly. Recently, LecRK-VI.2 was shown to interact with the immune co-receptor BAK1 (Huang *et al.,* 2014; Wang *et al.,* 2019) and both proteins were found to be necessary for eNAD^+^- and eNADP^+^-induced signaling. Given the critical nature of such interactions for signal transduction, solvent-exposed natural variants associated with reduced HR-like symptoms could affect the binding of putative co-receptors and partners of LecRK-I.1. Moreover, LecRK homo- and heterodimerization was reported (Bellande *et al.,* 2017; Wang & Bouwmeester, 2017), suggesting that a similar process might underlie LecRK-I.1 function and such interaction may be affected in the natural variants identified.

At the genomic level, *L-type LecRKs* are organized in clusters of tandem repeats that arose through local and whole genome duplication events (Hofberger *et al.,* 2015). Clade I *L-type LecRK* genes appear to be mostly Brassicaceae-specific (Hofberger *et al.,* 2015; Wang & Bouwmeester, 2017) and most of them have been involved in immunity. In particular, clade I *L-type LecRKs* originated following the At-α whole genome duplication event (50 Mya), and LecRKs that were duplicated during this period show signs of positive selection (Hofberger *et al.,* 2015). Consistent with this study, we found that the two major *LecRK-I.1* haplotypes segregate at intermediate to high frequencies, independently of any obvious geographical or phylogenetical pattern. The co-occurrence of such diverged polymorphisms indicate that they could be ancient and may have been maintained by selection. Consistent with the latter hypothesis, we found that the *LecRK-I.1* locus does not seem to evolve neutrally, as indicated by positive Tajima’s D as well as Fu and Li’s D/F statistics. Positive values for these statistics are considered as signatures of balancing selection, a type of selection that is observed in immunity-related or resistance genes for instance (Bakker *et al.,* 2006; Vila-Aiub *et al.,* 2011; Huard-Chauveau *et al.,* 2013; van Velzen & Etienne, 2015; Ariga *et al.,* 2017). Balancing selection collectively refers to processes by which genetic variation is maintained in a population, as opposed to purifying and positive selection that reduces variation (Delph & Kelly, 2014; Wu *et al.,* 2017). Different selective processes can ultimately lead to the maintenance of genetic variation such as overdominance (or heterozygote advantage), frequency-dependent selection or environmental heterogeneity (spatially varying selection). Identification of the exact mode of balancing selection acting at this locus deserves further investigation. Nevertheless, the fact that *LecRK-I.1* displays signatures of balancing selection highlights the potential ecological importance of this gene in natural Arabidopsis populations.

While the results presented here allow to move forward in the mechanistic understanding of the induction of cell death response in Arabidopsis, it is not known whether this gene may also play a role in crop species such as cabbage or canola. Interestingly, induction of HR-like in *B. nigra* also correlates with higher egg parasitism, indicating that both defense mechanisms may be under a similar regulation (Fatouros *et al.,* 2014). Further study of this region in other Brassicaceae and a screen of available germplasms for existing variation may allow to demonstrate a role for clade I LecRKs in sister species.

## Acknowledgements

We would like to thank Esteban Alfonso for technical help and Blaise Tissot for maintenace of the plants. We thank Dany Buffat for his technical assistance during the screening. We are also grateful to Nicolas Salamin and Anna Marcionetti for advices regarding the selection analyses. This work was supported by the Swiss National Science Foundation (grant no 31003A_169278 to P.R.) and EMBO (long term fellowship ALTF 248-2018 to E.K.).

## Author Contribution

RG and PR conceived and designed the research. RG and CG-D performed experiments. RG, EK and PR analyzed the data. RG and PR and wrote the manuscript.

