## Supplementary Material for "Natural variation in insect egg-induced cell death uncovers a role for L-type LECTIN RECEPTOR KINASE-I.1 in Arabidopsis"

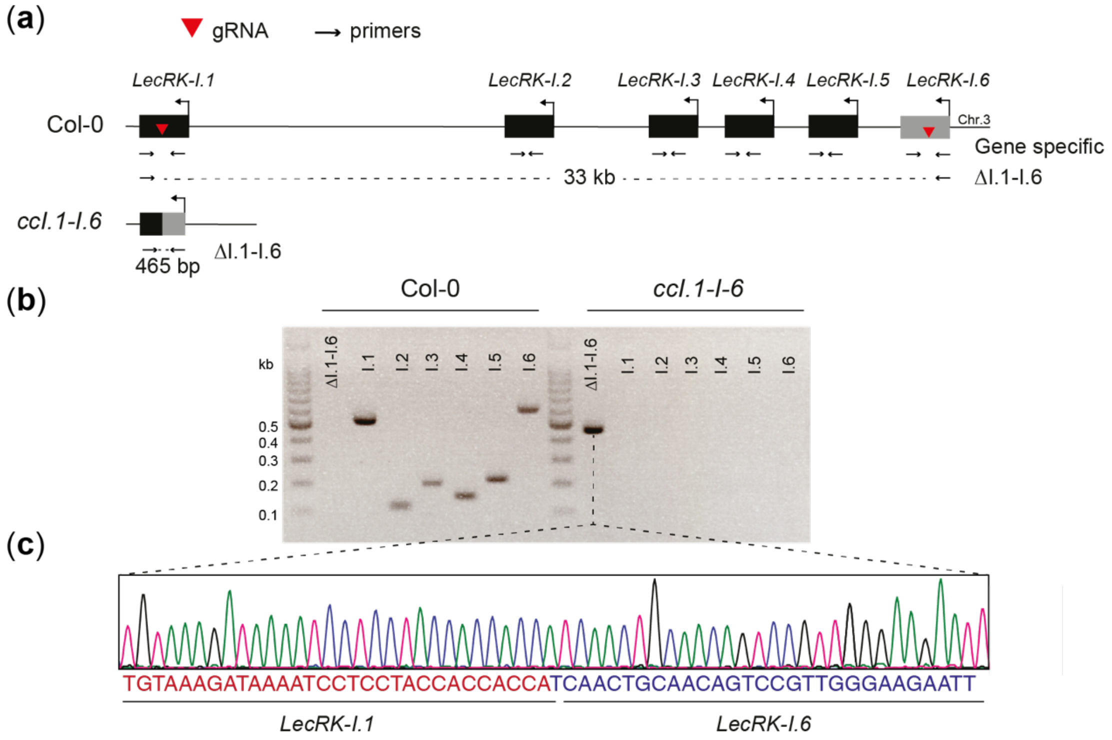


**Fig. S1** Deletion of *LecRK-I.1* to *LecRK-I.6* using CRISPR-Cas9. (a) Genomic cluster on chromosome 3 containing *LecRK-I.1* to *LecRK-I.6*. The two gRNA used to delete this six gene cluster are indicated with a red triangle. A simulation of the resulting chromosomal locus is depicted below. Arrows indicate primers used for genotyping and the size of the respective PCR products is shown. (b) PCR analysis of the *ccI.1-I.6* line using primers to identify the presence of a successful deletion (ΔI.1-I.6) and gene specific primers (I.1 to I.6). (c) The PCR product using the deletion-specific primers was sequenced and blasted against the Arabidopsis genome. Blast results reveal that the 33Kb region was successfully deleted.

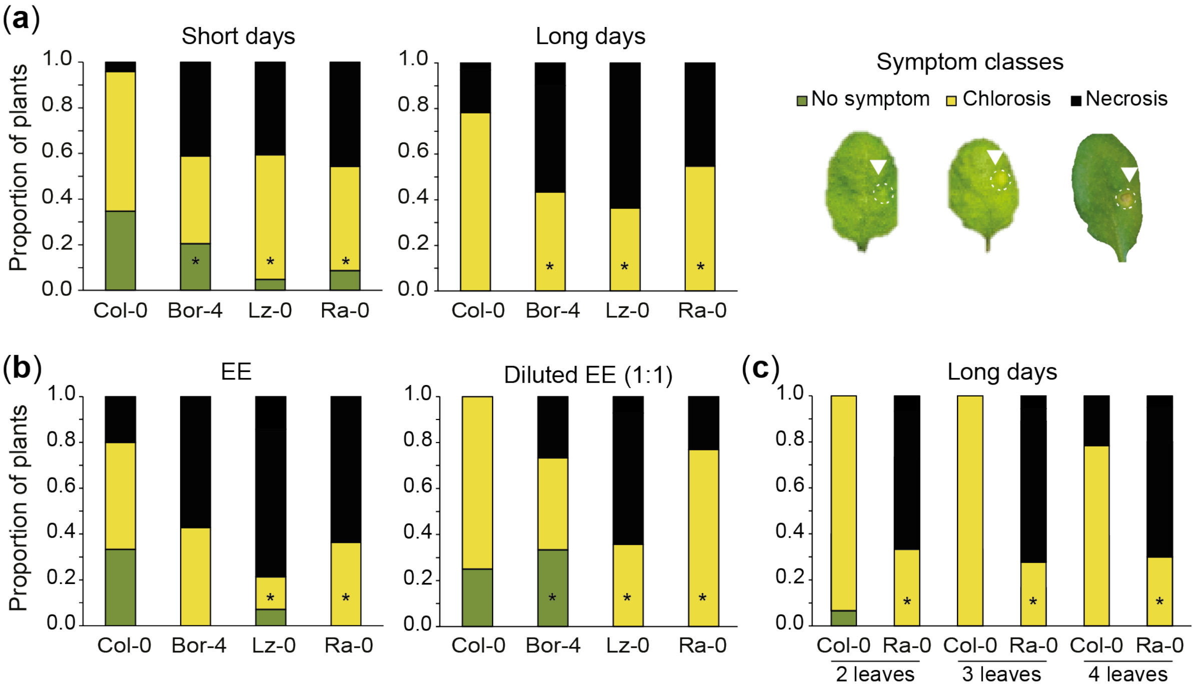


**Fig. S2** Setting-up conditions for the GWAS experiment. (a) Symptom distribution according to day length conditions after 5 days of EE treatment. Short days, 10 h light; Long days, 16 h light (b) Symptom distribution depending on whether EE was used intact or diluted 1:1 in distilled water. (c) Symptom distribution in Col-0 and Ra-0 after treatment of two, three or four leaves per plant. Unless specified, all experiments were carried out in short days condition and two leaves per plants were treated with EE. Symptoms were observed from the adaxial side of the leaves and classified according to the legend on the right. Asterisks denote significant differences between a given accession and Col-0 at $P$<0.05 (Fisher's exact test).

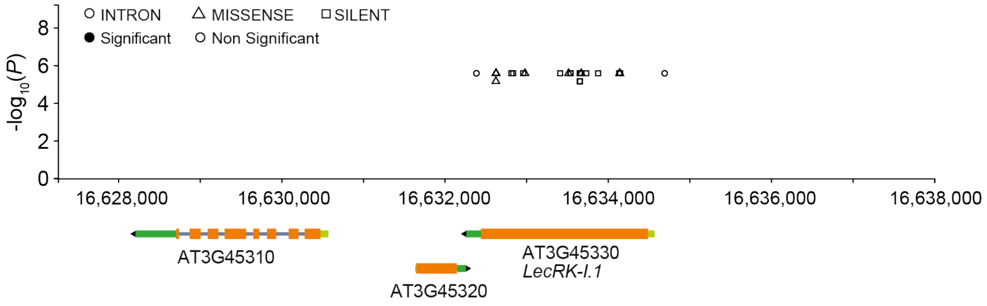


**Fig. S3** Meta-analysis of association at the *LecRK-I.1* locus. Screenshot of associations from the AraGWAS database at the *LecRK-I.1 locus*. X-axis represents genomic position along chromosome 3 and gene structures are shown below. Markers are shown depending on their genic position and on significance. Significance threshold was calculated based on permutations for each phenotype as described in Togninalli *et al.* (2018). None of the displayed markers reached significance. Only markers with MAF>5% are shown.

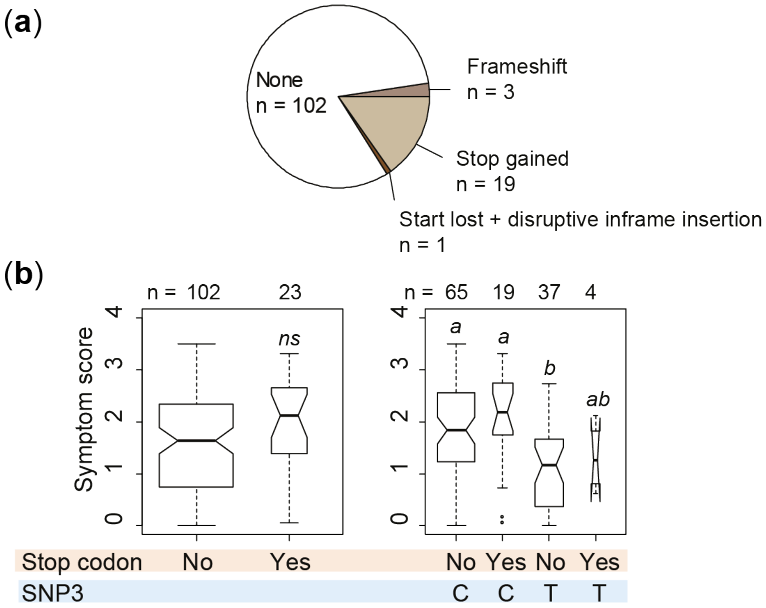


**Fig. S4** Disruptive variation in *LecRK-I.1* does not correlate with symptoms. (a) Proportion of sequenced accessions that contained premature stop codons and or frameshifts in their *LecRK-I.1* gene sequence. Different colors indicate the different types of disruptive polymorphisms found as annotated on POLYMORPH1001 (https://tools.1001genomes.org/polymorph/). N indicates the number of accessions that possessed a given type of variation. (b) HR-like symptom score distribution depending on the presence of disruptive variation (left panel) and evaluation of the effect of this variation depending on the allele present at SNP3. Different letters indicate significant difference at $P<0.05$ (Two- way ANOVA, followed by Tukey's HSD for multiple comparison). ns, not significant.

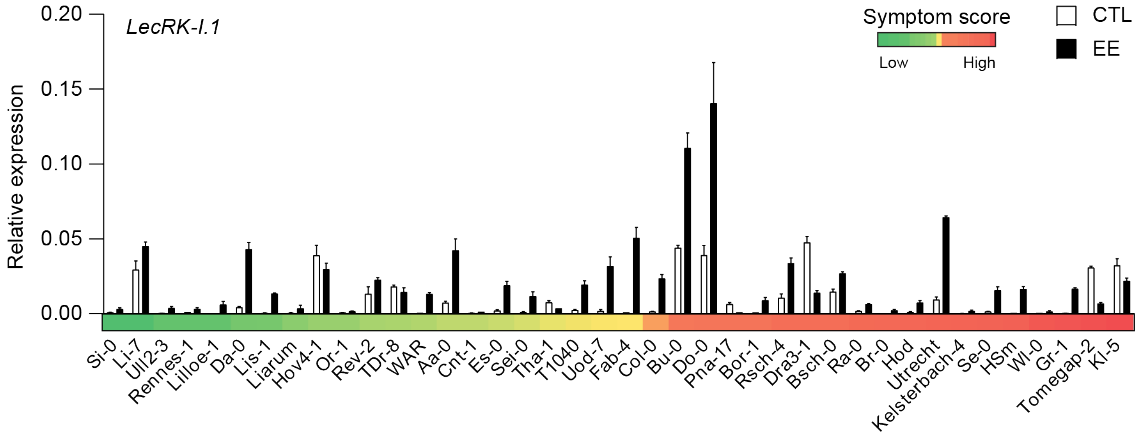


**Fig. S5** *LecRK-I.1* expression in a panel of 40 accessions with low or high symptom scores Expression of *LecRK-I.1* in 40 different accessions after 72 h of EE treatment. Gene expression was normalized to the reference gene *SAND*. Means ± SE of three technical replicates are shown. The average symptom score for each accession is shown using a color scale for clarity.

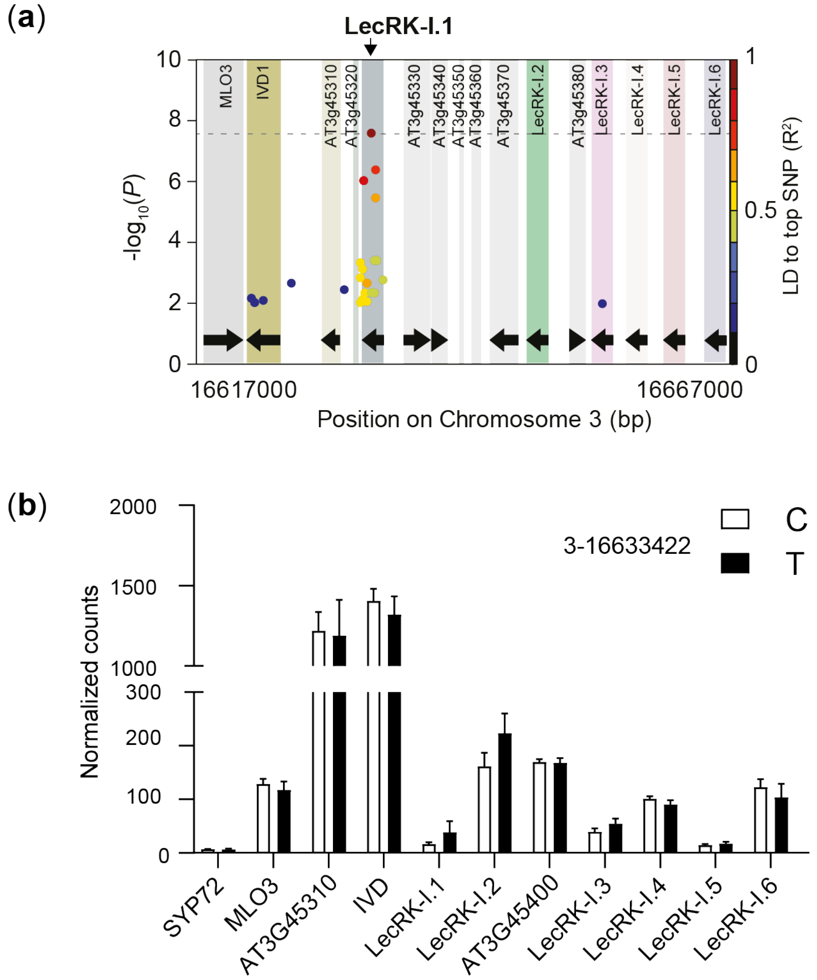


**Fig. S6** Basal expression of genes in the vicinity of the *LecRK-I.1* locus. (a,b) Publically available RNA-seq data were obtained for part of the accessions used in this study (54 out of 295) and expression levels for all genes in a 50 kb region around *LecRK-I.1* is shown depending on the allele present at SNP3 (C, n=42; T, n=12).

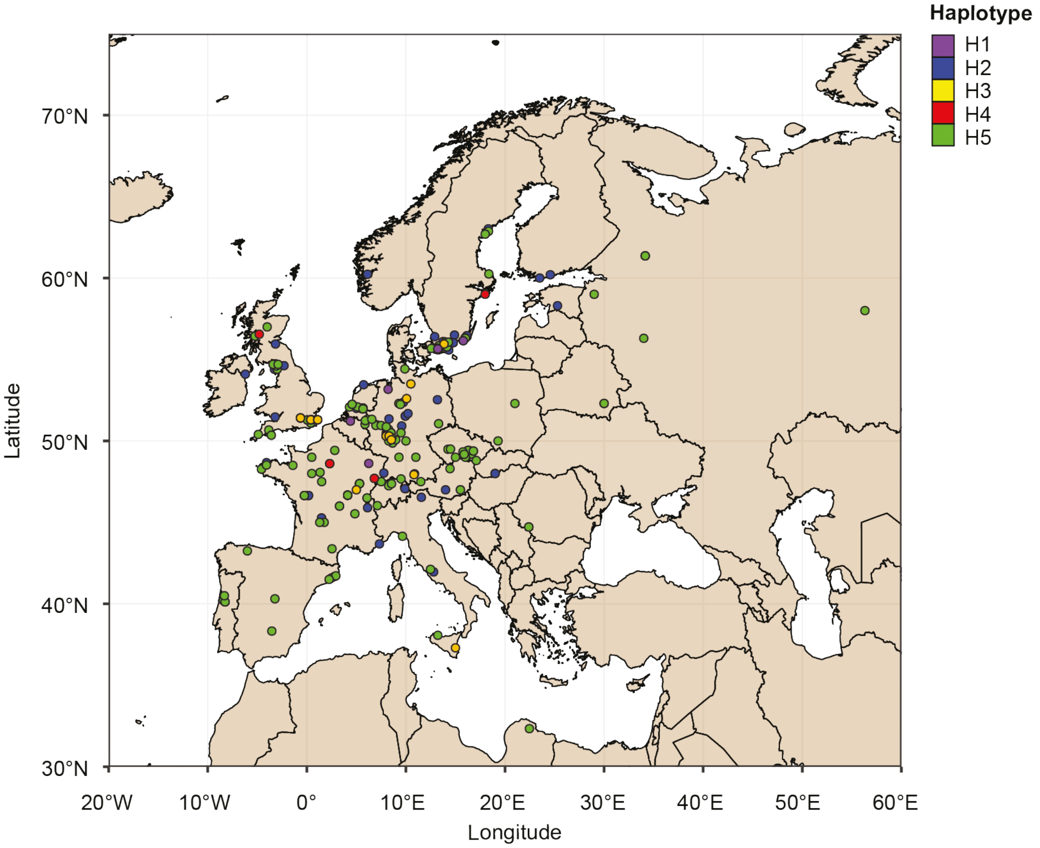


**Fig. S7** Map of *LecRK-I.1* haplotype distribution in European Arabidopsis accessions. Colors indicate haplotype identity. Out of the total 295 accessions used in this study, 20 originate from outside Europe and are not shown on this map for sake of clarity.

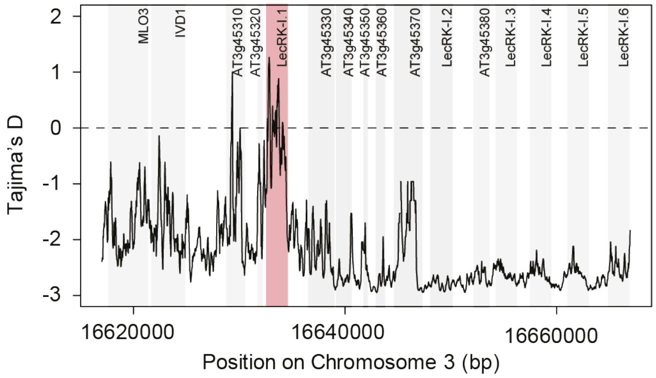


**Fig. S8** Sliding window analysis of Tajima's D in a 50kb region around *LecRK-I.1*. Tajima's D was computed using a window size of 200 bp and a step size of 25 bp. Rectangles indicate genes. A subset of 125 accessions with available full genome sequences was used for this analysis.

**Table S1.** List of the 295 Arabidopsis accessions used in this study. Name, accession ID and NASC line number are provided. Longitude and Latitude refer to geographical coordinates of the collection site, while "sequenced" indicates whether a given accession was sequenced as part of the 1001 Genomes project.

| Accession name | Accession ID | NASC number | Country | Latitude | Longitude | Sequenced? |
|---|---|---|---|---|---|---|
| ALL1-2 | 1 | N76089 | FRA | 45.2667 | 1.48333 | |
| ALL1-3 | 2 | N76090 | FRA | 45.2667 | 1.48333 | |
| CAM-16 | 23 | N76107 | FRA | 48.2667 | -4.58333 | |
| CAM-61 | 66 | N76108 | FRA | 48.2667 | -4.58333 | |
| CUR-3 | 81 | N76115 | FRA | 45 | 1.75 | |
| JEA | 91 | N76148 | FRA | 43.6833 | 7.33333 | |
| LAC-3 | 94 | N76157 | FRA | 47.7 | 6.81667 | |
| LAC-5 | 96 | N76158 | FRA | 47.7 | 6.81667 | |
| LDV-58 | 149 | N76163 | FRA | 48.5167 | -4.06667 | |
| MIB-15 | 166 | N76181 | FRA | 47.3833 | 5.31667 | |
| MIB-22 | 173 | N76182 | FRA | 47.3833 | 5.31667 | |
| MIB-28 | 178 | N76183 | FRA | 47.3833 | 5.31667 | |
| MIB-84 | 223 | N76184 | FRA | 47.3833 | 5.31667 | |
| MOG-37 | 242 | N76189 | FRA | 48.6667 | -4.06667 | |
| PAR-3 | 258 | N76205 | FRA | 46.65 | -0.25 | |
| PAR-4 | 259 | N76206 | FRA | 46.65 | -0.25 | |
| PAR-5 | 260 | N76207 | FRA | 46.65 | -0.25 | |
| ROM-1 | 267 | N76221 | FRA | 45.5333 | 4.85 | |
| TOU-A1-115 | 281 | N76252 | FRA | 46.6667 | 4.11667 | |
| TOU-A1-116 | 282 | N76253 | FRA | 46.6667 | 4.11667 | |
| TOU-A1-43 | 321 | N76255 | FRA | 46.6667 | 4.11667 | |
| TOU-A1-62 | 328 | N76256 | FRA | 46.6667 | 4.11667 | |
| TOU-A1-96 | 357 | N76258 | FRA | 46.6667 | 4.11667 | |
| TOU-C-3 | 362 | N76259 | FRA | 46.6667 | 4.11667 | |
| TOU-E-11 | 366 | N76260 | FRA | 46.6667 | 4.11667 | |
| TOU-H-12 | 373 | N76261 | FRA | 46.6667 | 4.11667 | |
| TOU-H-13 | 374 | N76262 | FRA | 46.6667 | 4.11667 | |
| TOU-I-17 | 378 | N76263 | FRA | 46.6667 | 4.11667 | |
| TOU-I-6 | 380 | N76265 | FRA | 46.6667 | 4.11667 | |
| TOU-J-3 | 383 | N76266 | FRA | 46.6667 | 4.11667 | |
| VOU-1 | 390 | N76299 | FRA | 46.65 | 0.166667 | |
| VOU-2 | 392 | N76300 | FRA | 46.65 | 0.166667 | |
| LI-OF-095 | 641 | N76165 | USA | 40.7777 | -72.9069 | |
| Belmonte-4-94 | 957 | N76095 | ITA | 42.1167 | 12.4833 | |
| KBS-Mac-8 | 1716 | N76151 | USA | 42.405 | -85.398 | |
| MNF-Pot-48 | 1859 | N76187 | USA | 43.595 | -86.2657 | |
| MNF-Pot-68 | 1867 | N76188 | USA | 43.595 | -86.2657 | |
| MNF-Jac-32 | 1967 | N76186 | USA | 43.5187 | -86.1739 | |
| Map-42 | 2057 | N76180 | USA | 42.166 | -86.412 | Yes |
| Paw-3 | 2150 | N76208 | USA | 42.148 | -86.431 | |
| Pent-1 | 2187 | N76209 | USA | 43.7623 | -86.3929 | |
| SLSP-30 | 2274 | N76228 | USA | 43.665 | -86.496 | |
| Ste-3 | 2290 | N76232 | USA | 42.03 | -86.514 | |
| UKSW06-202 | 4802 | N76292 | UK | 50.4 | -4.9 | |
| UKSE06-062 | 4997 | N76280 | UK | 51.3 | 0.5 | |
| UKSE06-192 | 5056 | N76281 | UK | 51.3 | 0.5 | |
| UKSE06-272 | 5116 | N76282 | UK | 51.3 | 0.4 | |

| Accession name | Accession ID | NASC number | Country | Latitude | Longitude | Sequenced? |
|---|---|---|---|---|---|---|
| UKSE06-349 | 5158 | N76284 | UK | 51.3 | 0.4 | |
| UKSE06-351 | 5160 | N76285 | UK | 51.3 | 0.4 | |
| UKSE06-414 | 5202 | N76286 | UK | 51.3 | 0.4 | |
| UKSE06-429 | 5207 | N76287 | UK | 51.3 | 0.4 | |
| UKSE06-466 | 5232 | N76288 | UK | 51.2 | 0.4 | |
| UKSE06-520 | 5264 | N76290 | UK | 51.3 | 1.1 | |
| UKSE06-628 | 5341 | N76291 | UK | 51.1 | 0.4 | |
| UKNW06-059 | 5380 | N76275 | UK | 54.4 | -3 | |
| UKNW06-060 | 5381 | N76276 | UK | 54.4 | -3 | |
| UKNW06-386 | 5565 | N76277 | UK | 54.6 | -3.1 | |
| UKNW06-436 | 5606 | N76278 | UK | 54.7 | -3.4 | |
| UKNW06-460 | 5628 | N76279 | UK | 54.7 | -3.4 | |
| Bur-0 | 5719 | N76105 | IRL | 53.08 | -9.0755 | |
| UKID22 | 5729 | N76271 | UK | 54.7 | -3.4 | |
| UKID37 | 5742 | N76272 | UK | 51.3 | 1.1 | |
| UKID80 | 5785 | N76274 | UK | 54.7 | -2.9 | |
| App1-16 | 5832 | N76092 | SWE | 56.3333 | 15.9667 | Yes |
| Bor-1 | 5837 | N76099 | CZE | 49.4013 | 16.2326 | Yes |
| DraIV1-14 | 5896 | N76119 | CZE | 49.4112 | 16.2815 | |
| DraIV6-16 | 5987 | N76122 | CZE | 49.4112 | 16.2815 | |
| DraIV6-35 | 6005 | N76123 | CZE | 49.4112 | 16.2815 | |
| Duk | 6008 | N76124 | CZE | 49.1 | 16.2 | Yes |
| Fja1-2 | 6019 | N76131 | SWE | 56.06 | 14.29 | Yes |
| Fja1-5 | 6020 | N76132 | SWE | 56.06 | 14.29 | Yes |
| Lom1-1 | 6042 | N76174 | SWE | 56.09 | 13.9 | Yes |
| Lov-5 | 6046 | N76175 | SWE | 62.801 | 18.079 | Yes |
| Or-1 | 6074 | N76201 | SWE | 56.4573 | 16.1308 | Yes |
| Rev-2 | 6076 | N76219 | SWE | 55.6942 | 13.4504 | Yes |
| Sparta-1 | 6085 | N76229 | SWE | 55.7097 | 13.2145 | Yes |
| T1040 | 6094 | N76233 | SWE | 55.6494 | 13.2147 | Yes |
| T1060 | 6096 | N76234 | SWE | 55.6472 | 13.2225 | Yes |
| T1080 | 6098 | N76235 | SWE | 55.6561 | 13.2178 | Yes |
| T1110 | 6100 | N76236 | SWE | 55.6 | 13.2 | Yes |
| T1130 | 6102 | N76237 | SWE | 55.6 | 13.2 | Yes |
| T690 | 6124 | N76241 | SWE | 55.8378 | 13.3092 | Yes |
| Tad01 | 6169 | N76243 | SWE | 62.8714 | 18.3447 | Yes |
| TDr-1 | 6188 | N76245 | SWE | 55.7683 | 14.1386 | Yes |
| TDr-3 | 6190 | N76248 | SWE | 55.7686 | 14.1381 | |
| TDr-8 | 6194 | N76249 | SWE | 55.7706 | 14.1342 | Yes |
| TDr-18 | 6203 | N76247 | SWE | 55.7714 | 14.1208 | Yes |
| Tomegap-2 | 6242 | N76250 | SWE | 55.7 | 13.2 | Yes |
| Tottarp-2 | 6243 | N76251 | SWE | 56.27373 | 13.90045 | Yes |
| Udul1-34 | 6318 | N76269 | CZE | 49.2771 | 16.6314 | |
| Ull3-4 | 6413 | N76295 | SWE | 56.06 | 13.97 | Yes |
| Zdrl2-25 | 6449 | N76308 | CZE | 49.3853 | 16.2544 | |
| Bg2 | 6709 | N76096 | USA | 47.6479 | -122.305 | |
| CIBC2 | 6727 | N28140 | UK | 51.4083 | -0.6383 | |
| CIBC5 | 6730 | N28142 | UK | 51.4083 | -0.6383 | |
| Cold Spring Harbor Lab-5 | 6744 | N28181 | USA | 40.8585 | -73.4675 | Yes |
| NFC20 | 6847 | N28550 | UK | 51.4083 | -0.6383 | |
| Ag-0 | 6897 | N76087 | FRA | 45 | 1.3 | Yes |
| An-1 | 6898 | N76091 | BEL | 51.2167 | 4.4 | Yes |
| Bay-0 | 6899 | N76094 | GER | 49 | 11 | |

| Accession name | Accession ID | NASC number | Country | Latitude | Longitude | Sequenced? |
|---|---|---|---|---|---|---|
| Bor-4 | 6903 | N76100 | CZE | 49.4013 | 16.2326 | Yes |
| Br-0 | 6904 | N76101 | CZE | 49.2 | 16.6166 | Yes |
| C24 | 6906 | N76106 | POR | 40.2077 | -8.42639 | |
| CIBC17 | 6907 | N76111 | UK | 51.4083 | -0.6383 | Yes |
| Col-0 | 6909 | N76113 | USA | 38.3 | -92.3 | Yes |
| Ct-1 | 6910 | N76114 | ITA | 37.3 | 15 | |
| Cvi-0 | 6911 | N76116 | CPV | 15.1111 | -23.6167 | Yes |
| Edi-0 | 6914 | N76126 | UK | 55.9494 | -3.16028 | |
| Fab-4 | 6918 | N76128 | SWE | 63.0165 | 18.3174 | Yes |
| Ga-0 | 6919 | N76133 | GER | 50.3 | 8 | Yes |
| Got-7 | 6921 | N76136 | GER | 51.5338 | 9.9355 | |
| HR-5 | 6924 | N76144 | UK | 51.4083 | -0.6383 | Yes |
| Kin-0 | 6926 | N76153 | USA | 44.46 | -85.37 | Yes |
| Kno-18 | 6928 | N76154 | USA | 41.2816 | -86.621 | |
| Ler-1 | 6932 | N76164 | GER | 47.984 | 10.8719 | Yes |
| LL-0 | 6933 | N76172 | ESP | 41.59 | 2.49 | Yes |
| Lz-0 | 6936 | N76179 | FRA | 46 | 3.3 | |
| Mrk-0 | 6937 | N76191 | GER | 49 | 9.3 | |
| Mt-0 | 6939 | N76192 | LIB | 32.34 | 22.46 | |
| Mz-0 | 6940 | N76193 | GER | 50.3 | 8.3 | Yes |
| Nd-1 | 6942 | N76197 | GER | 50 | 10 | |
| NFA-10 | 6943 | N76198 | UK | 51.4083 | -0.6383 | Yes |
| Oy-0 | 6946 | N76203 | NOR | 60.39 | 6.19 | |
| Pu2-23 | 6951 | N76215 | CZE | 49.42 | 16.36 | Yes |
| Ra-0 | 6958 | N76216 | FRA | 46 | 3.3 | Yes |
| Rennes-1 | 6959 | N76218 | FRA | 48.5 | -1.41 | Yes |
| Se-0 | 6961 | N76226 | ESP | 38.3333 | -3.53333 | Yes |
| Sha | 6962 | N76227 | TJK | 38.35 | 68.48 | |
| Tamm-2 | 6968 | N76244 | FIN | 60 | 23.5 | Yes |
| Ts-1 | 6970 | N76268 | ESP | 41.7194 | 2.93056 | Yes |
| Ull2-3 | 6973 | N76293 | SWE | 56.0648 | 13.9707 | Yes |
| Ull2-5 | 6974 | N76294 | SWE | 56.0648 | 13.9707 | Yes |
| Uod-7 | 6976 | N76296 | AUT | 48.3 | 14.45 | Yes |
| Van-0 | 6977 | N76297 | CAN | 49.3 | -123 | |
| Wa-1 | 6978 | N28804 | POL | 52.3 | 21 | |
| Wei-0 | 6979 | N76301 | SUI | 47.25 | 8.26 | Yes |
| Ws-0 | 6980 | N76303 | RUS | 52.3 | 30 | |
| Wt-5 | 6982 | N76304 | GER | 52.3 | 9.3 | Yes |
| Yo-0 | 6983 | N76305 | USA | 37.45 | -119.35 | |
| Zdr-6 | 6985 | N76306 | CZE | 49.3853 | 16.2544 | |
| Alc-0 | 6988 | N76088 | ESP | 40.49 | -3.36 | |
| Amel-1 | 6990 | N28014 | NED | 53.448 | 5.73 | Yes |
| Ann-1 | 6994 | N28049 | FRA | 45.9 | 6.13028 | |
| An-2 | 6996 | N28017 | BEL | 51.2167 | 4.4 | |
| Aa-0 | 7000 | N28007 | GER | 50.9167 | 9.57073 | Yes |
| Baa-1 | 7002 | N28054 | NED | 51.3333 | 6.1 | Yes |
| Bs-2 | 7004 | N28097 | SUI | 47.5 | 7.5 | |
| Benk-1 | 7008 | N28064 | NED | 52 | 5.675 | Yes |
| Ba-1 | 7014 | N28053 | UK | 56.5459 | -4.79821 | Yes |
| Bla-1 | 7015 | N76097 | ESP | 41.6833 | 2.8 | |
| Boot-1 | 7026 | N28091 | UK | 54.4 | -3.2667 | Yes |
| Bsch-0 | 7031 | N28099 | GER | 50.0167 | 8.6667 | Yes |
| Blh-1 | 7034 | N76098 | CZE | 48.3 | 19.85 | |
| Blh-2 | 7035 | N28090 | CZE | 48 | 19 | |

| Accession name | Accession ID | NASC number | Country | Latitude | Longitude | Sequenced? |
|---|---|---|---|---|---|---|
| Ca-0 | 7062 | N28128 | GER | 50.2981 | 8.26607 | Yes |
| Cnt-1 | 7064 | N28160 | UK | 51.3 | 1.1 | Yes |
| Cha-0 | 7069 | N28133 | SUI | 46.0333 | 7.1167 | |
| Chat-1 | 7071 | N28135 | FRA | 48.0717 | 1.33867 | Yes |
| Cit-0 | 7075 | N28158 | FRA | 43.3779 | 2.54038 | |
| Co-2 | 7078 | N28163 | POR | 40.12 | -8.25 | |
| Co-4 | 7080 | N28165 | POR | 40.12 | -8.25 | |
| Com-1 | 7092 | N28193 | FRA | 49.416 | 2.823 | Yes |
| Da-0 | 7094 | N28200 | GER | 49.8724 | 8.65081 | Yes |
| Di-1 | 7098 | N28208 | FRA | 47 | 5 | |
| Db-0 | 7100 | N28202 | GER | 50.3055 | 8.324 | |
| Do-0 | 7102 | N28210 | GER | 50.7224 | 8.2372 | Yes |
| Dra-2 | 7105 | N28214 | CZE | 49.4167 | 16.2667 | |
| Ede-1 | 7110 | N28217 | NED | 52.0333 | 5.66667 | |
| Ep-0 | 7123 | N28236 | GER | 50.1721 | 8.38912 | |
| Es-0 | 7126 | N28241 | FIN | 60.1997 | 24.5682 | Yes |
| Est-0 | 7128 | N28243 | RUS | 58.3 | 25.3 | |
| Fr-4 | 7135 | N28268 | GER | 50.1102 | 8.6822 | |
| Fi-1 | 7139 | N28252 | GER | 50.5 | 8.0167 | |
| Ga-2 | 7141 | N28274 | GER | 50.3 | 8 | |
| Gel-1 | 7143 | N28279 | NED | 51.0167 | 5.86667 | Yes |
| Ge-1 | 7145 | N28277 | SUI | 46.5 | 6.08 | |
| Gü• -1 | 7150 | N28332 | GER | 50.3 | 8 | |
| Gö-0 | 7151 | N28282 | GER | 51.5338 | 9.9355 | |
| Gr-5 | 7158 | N28326 | AUT | 47 | 15.5 | Yes |
| Ha-0 | 7163 | N28336 | GER | 52.3721 | 9.73569 | Yes |
| Hau-0 | 7164 | N28343 | DEN | 55.675 | 12.5686 | Yes |
| Hn-0 | 7165 | N28350 | GER | 51.3472 | 8.28844 | Yes |
| Hey-1 | 7166 | N28344 | NED | 51.25 | 5.9 | |
| Hh-0 | 7169 | N28345 | GER | 54.4175 | 9.88682 | Yes |
| Jm-1 | 7178 | N28373 | CZE | 49 | 15 | |
| Kelsterbach -2 | 7188 | N28382 | GER | 50.0667 | 8.5333 | |
| Kl-5 | 7199 | N28394 | GER | 50.95 | 6.9666 | Yes |
| Kr-0 | 7201 | N28419 | GER | 51.3317 | 6.55934 | |
| Kro-0 | 7206 | N28420 | GER | 50.0742 | 8.96617 | |
| Li-3 | 7224 | N28454 | GER | 50.3833 | 8.0666 | |
| Li-5:2 | 7227 | N28457 | GER | 50.3833 | 8.0666 | |
| Li-6 | 7229 | N28459 | GER | 50.3833 | 8.0666 | |
| Li-7 | 7231 | N28461 | GER | 50.3833 | 8.0666 | Yes |
| Mnz-0 | 7244 | N28495 | GER | 50.001 | 8.26664 | Yes |
| Mc-0 | 7252 | N28490 | UK | 54.6167 | -2.3 | |
| Mh-0 | 7255 | N28492 | POL | 50.95 | 20.5 | Yes |
| Nw-0 | 7258 | N28573 | GER | 50.5 | 8.5 | Yes |
| Nw-2 | 7260 | N28575 | GER | 50.5 | 8.5 | |
| Nz1 | 7263 | N28578 | NZL | -37.7871 | 175.283 | |
| No-0 | 7275 | N28564 | GER | 51.0581 | 13.2995 | |
| Ob-1 | 7277 | N28580 | GER | 50.2 | 8.5833 | |
| Old-1 | 7280 | N28583 | GER | 53.1667 | 8.2 | Yes |
| Or-0 | 7282 | N28587 | GER | 50.3827 | 8.01161 | Yes |
| Ors-1 | 7283 | N28848 | ROU | 44.7203 | 22.3955 | |
| Ors-2 | 7284 | N28849 | ROU | 44.7203 | 22.3955 | |
| Petergof | 7296 | N76211 | RUS | 59 | 29 | Yes |
| Pla-0 | 7300 | N28640 | ESP | 41.5 | 2.25 | |
| Rhen-1 | 7316 | N28685 | NED | 51.9667 | 5.56667 | Yes |

| Accession name | Accession ID | NASC number | Country | Latitude | Longitude | Sequenced? |
|---|---|---|---|---|---|---|
| Sh-0 | 7331 | N28734 | GER | 51.6832 | 10.2144 | |
| Sei-0 | 7333 | N28729 | ITA | 46.5438 | 11.5614 | Yes |
| Si-0 | 7337 | N28739 | GER | 50.8738 | 8.02341 | Yes |
| Sp-0 | 7343 | N28743 | GER | 52.5339 | 13.181 | Yes |
| Sg-1 | 7344 | N28732 | GER | 47.6667 | 9.5 | Yes |
| Ty-0 | 7351 | N28786 | UK | 56.4278 | -5.23439 | |
| Tha-1 | 7353 | N28758 | NED | 52.08 | 4.3 | Yes |
| Ting-1 | 7354 | N28759 | SWE | 56.5 | 14.9 | Yes |
| Tiv-1 | 7355 | N28760 | ITA | 41.96 | 12.8 | |
| Tscha-1 | 7372 | N28779 | AUT | 47.0748 | 9.9042 | Yes |
| Tsu-0 | 7373 | N28780 | JPN | 34.43 | 136.31 | Yes |
| Uk-1 | 7378 | N28787 | GER | 48.0333 | 7.7667 | Yes |
| Utrecht | 7382 | N28795 | NED | 52.0918 | 5.1145 | Yes |
| Wag-3 | 7390 | N28808 | NED | 51.9666 | 5.6666 | |
| Wag-4 | 7391 | N28809 | NED | 51.9666 | 5.6666 | |
| Wag-5 | 7392 | N28810 | NED | 51.9666 | 5.6666 | |
| Ws | 7397 | N28823 | RUS | 52.3 | 30 | |
| Wc-2 | 7405 | N28814 | GER | 52.6 | 10.0667 | |
| Wt-3 | 7408 | N28833 | GER | 52.3 | 9.3 | |
| Wl-0 | 7411 | N28822 | GER | 47.9299 | 10.8134 | Yes |
| Zu-1 | 7418 | N28847 | SUI | 47.3667 | 8.55 | Yes |
| Nc-1 | 7430 | N28527 | FRA | 48.6167 | 6.25 | Yes |
| N13 | 7438 | N76194 | RUS | 61.36 | 34.15 | |
| N4 | 7446 | N28510 | RUS | 61.36 | 34.15 | |
| N7 | 7449 | N28513 | RUS | 61.36 | 34.15 | |
| WAR | 7477 | N28812 | USA | 41.7302 | -71.2825 | Yes |
| PHW-10 | 7479 | N28610 | UK | 51.2878 | 0.0565 | |
| PHW-13 | 7482 | N28613 | UK | 51.2878 | 0.0565 | |
| PHW-20 | 7490 | N28620 | UK | 51.2878 | 0.0565 | |
| PHW-26 | 7496 | N28626 | UK | 50.6728 | -3.8404 | |
| PHW-28 | 7498 | N28628 | UK | 50.35 | -3.5833 | |
| PHW-31 | 7502 | N28631 | UK | 51.4666 | -3.2 | |
| PHW-33 | 7504 | N28633 | NED | 52.25 | 4.5667 | |
| PHW-36 | 7507 | N28636 | FRA | 48.6103 | 2.3086 | |
| PHW-37 | 7508 | N28637 | FRA | 48.6103 | 2.3086 | |
| Var2-1 | 7516 | N76298 | SWE | 55.58 | 14.334 | Yes |
| Omo2-1 | 7518 | N76200 | SWE | 56.14 | 15.78 | |
| Lp2-2 | 7520 | N76176 | CZE | 49.38 | 16.81 | Yes |
| Lp2-6 | 7521 | N76177 | CZE | 49.38 | 16.81 | Yes |
| Mr-0 | 7522 | N76190 | ITA | 44.15 | 9.65 | |
| Pna-17 | 7523 | N76213 | USA | 42.0945 | -86.3253 | Yes |
| Rmx-A180 | 7525 | N76220 | USA | 42.036 | -86.511 | Yes |
| Pro-0 | 8213 | N76214 | ESP | 43.25 | -6 | |
| Fei-0 | 8215 | N76129 | POR | 40.5 | -8.32 | |
| Lis-2 | 8222 | N76170 | SWE | 56.0328 | 14.775 | Yes |
| Bro1-6 | 8231 | N76102 | SWE | 56.3 | 16 | Yes |
| Hod | 8235 | N76141 | CZE | 48.8 | 17.1 | Yes |
| HSm | 8236 | N76146 | CZE | 49.33 | 15.76 | Yes |
| PHW-3 | 8239 | N76155 | GER | 51 | 7 | Yes |
| Liarum | 8241 | N76166 | SWE | 55.9473 | 13.821 | Yes |
| Lillo-1 | 8242 | N76167 | SWE | 56.1494 | 15.7884 | Yes |
| Ba1-2 | 8256 | N76093 | SWE | 56.4 | 12.9 | Yes |
| Bu-0 | 8271 | N76103 | GER | 50.5 | 9.5 | |
| Can-0 | 8274 | N76109 | ESP | 29.2144 | -13.4811 | |

| Accession name | Accession ID | NASC number | Country | Latitude | Longitude | Sequenced? |
|---|---|---|---|---|---|---|
| Cen-0 | 8275 | N76110 | FRA | 49 | 0.5 | |
| Dra3-1 | 8283 | N76117 | SWE | 55.76 | 14.12 | Yes |
| Gd-1 | 8296 | N76134 | GER | 53.5 | 10.5 | |
| Ge-0 | 8297 | N76135 | SUI | 46.5 | 6.08 | Yes |
| Gr-1 | 8300 | N76137 | AUT | 47 | 15.5 | |
| Hi-0 | 8304 | N76140 | NED | 52 | 5 | |
| Hov4-1 | 8306 | N76142 | SWE | 56.1 | 13.74 | Yes |
| Hs-0 | 8310 | N76145 | GER | 52.24 | 9.44 | |
| In-0 | 8311 | N76147 | AUT | 47.5 | 11.5 | Yes |
| Ka-0 | 8314 | N76149 | AUT | 47 | 14 | |
| Lc-0 | 8323 | N76159 | UK | 57 | -4 | |
| Lip-0 | 8325 | N76168 | POL | 50 | 19.3 | |
| Lis-1 | 8326 | N76169 | SWE | 56.0328 | 14.775 | Yes |
| Lm-2 | 8329 | N76173 | FRA | 48 | 0.5 | |
| Lund | 8335 | N76178 | SWE | 55.71 | 13.2 | Yes |
| Na-1 | 8343 | N76195 | FRA | 47.5 | 1.5 | Yes |
| Ost-0 | 8351 | N76202 | SWE | 60.25 | 18.37 | Yes |
| Pa-1 | 8353 | N76204 | ITA | 38.07 | 13.22 | |
| Per-1 | 8354 | N76210 | RUS | 58 | 56.3167 | Yes |
| Rak-2 | 8365 | N76217 | CZE | 49 | 16 | Yes |
| Rsch-4 | 8374 | N76222 | RUS | 56.3 | 34 | |
| Sanna-2 | 8376 | N76223 | SWE | 62.69 | 18 | Yes |
| Sap-0 | 8378 | N76224 | CZE | 49.49 | 14.24 | |
| St-0 | 8387 | N76231 | SWE | 59 | 18 | Yes |
| Ta-0 | 8389 | N76242 | CZE | 49.5 | 14.5 | |
| Sav-1 | 8412 | N76225 | CZE | 49.1833 | 15.8833 | |
| Kelsterbach -4 | 8420 | N76152 | GER | 50.0667 | 8.5333 | Yes |
| Fja1-1 | 8422 | N76130 | SWE | 56.06 | 14.29 | Yes |
| Kas-2 | 8424 | N76150 | IND | 35 | 77 | Yes |
| Lisse | 8430 | N76171 | NED | 52.25 | 4.5667 | |
| 11ME1.32 | 8610 | N76083 | USA | 42.093 | -86.359 | |
| 328PNA054 | 8692 | N76085 | USA | 42.0945 | -86.3253 | |
| 11PNA4.101 | 8796 | N76084 | USA | 42.0945 | -86.3253 | |

**Table S2** List of primers used in this study.

| Gene | Id | Primer | Sequence (5'-3') |
|---|---|---|---|
| QPCR | | | |
| *SAND* | At2g28390 | SAND_Fw | AACTCTATGCAGCATTTGATCCACT |
| | | SAND_Rv | TGATTGCATATCTTTATCGCCATC |
| *LecRK-I.1* | At3g45330 | LecRK-I.1_Fw | CCCGGATCGAAAAGCATTCA |
| | | LecRK-I.1_Rv | GTTTTCTCCGGTTTCTTGGG |
| | | LecRK-I.1_acc_FW* | GGTTTTCGGCTGCGACTGG |
| | | LecRK-I.1_acc_Rv* | AGATAAAATCCTCCTACCACC |
| *PR1* | At2g14610 | PR1_Fw | GTGGGTTAGCGAGAAGGCTA |
| | | PR1_Rv | ACTTTGGCACA TCCGAGTCT |
| CRISPR-Cas9 | | | |
| | | sgRNA_LecRK-I.1-Fw | **ATTG**ACTCGCTATCATAGTAATGG |
| | | sgRNA_LecRK-I.1-Rv | **AAAC**CCATTACTATGATAGCGAGT |
| | | sgRNA_LecRK-I.6-Fw | **ATTG**GACTGTTGCAGTTGACCGA |
| | | sgRNA_LecRK-I.6-Rv | **AAAC**TCGGTCAACTGCAACAGTC |
| Genotyping | | | |
| 33 kb Deletion | | CC-LecRK-I.6-Fw | GATCTGGGTGATCTTCTGTC |
| | | CC-LecRK-I.1-Rv | TTCGCCTACAATACCCGAGA |
| *LecRK-I.1* | At3g45330 | CC-LecRK-I.1-Fw | CCCGGATCGAAAAGCATTCA |
| | | CC-LecRK-I.1-Rv | TTCGCCTACAATACCCGAGA |
| *LecRK-I.2* | At3g45390 | LecRK-I.2_Fw | CCCGGATCGAAAAGCTTTTG |
| | | LecRK-I.2_Rv | GTTCAGAGATGTCAAGGCTC |
| *LecRK-I.3* | At3g45410 | LecRK-I.3_Fw | TCCTAAGTTGGGAGCTGATG |
| | | LecRK-I.3_Rv | ACCAACATGAGGCTTCTCCA |
| *LecRK-I.4* | At3g45420 | LecRK-I.4_Fw | TGAAACTTGGGTTGCTCTGC |
| | | LecRK-I.4_Rv | ATGGTTCCACTGAAACTGGC |
| *LecRK-I.5* | At3g45430 | LecRK-I.5_Fw | TGCTGAGTTGAACGGTAGGT |
| | | LecRK-I.5_Rv | CTCAACTTGCACCCCGAATT |
| *LecRK-I.6* | At3g45440 | CC-LecRK-I.6-Fw | GATCTGGGTGATCTTCTGTC |
| | | CC-LecRK-I.6-Rv | TTTAAGAGGGGCCAGCGTAA |

* primers used for *LecRK-I.1* expression analysis in different accessions (Fig. S5).
